# A mechanistic insight into *Nigella sativa* mediated anticancer effect on breast cancer through regulation of various miRNAs: An *in vitro* & *in vivo* study

**DOI:** 10.1101/2022.10.18.512670

**Authors:** Shaswati Das, Avijit Ghosh, Priyanka Upadhyay, Sushmita Sarker, Mousumi Bhattacharjee, Payal Gupta, Swatilekha Ghosh, Sreya Chattopadhyay, Pubali Dhar, Arghya Adhikary

**Author notes:** **Corresponding Author:** Dr. Arghya Adhikary. ^1^Centre for Research in Nanoscience and Nanotechnology, Technology Campus, University of Calcutta, JD-2, Sector III, Salt Lake City, Kolkata-700106, India.

## Abstract

**Background:** Cancer continues to threat the mortal alongside scientific community with its burgeoning grasp. Although efforts have been directed to tame cancer by radiotherapy, photodynamic therapy, chemotherapy it came at a cost of fatal side effects.

**Purpose:** Plant derived bioactive compounds carries an inevitable advantage of being safer, bioavailable & less toxic compared to contemporary chemotherapeutics. This study analyzed anti-cancerous potential of volatile oil, extracted from *Nigella sativa*, in-vitro against MDA-MB-231, MCF-7 & in-vivo on tumor growth in mice after successful oral administration.

**Study Design:** Our strategic approach employed solvent extraction of black seed oil (BSO) to highlight orchestrated use of its potent integrants - TQ, Carvacrol & TA which in modest amounts show anti-cancerous properties compared to their individual treatment.

We attempted to show this cost effective, safe & bioavailable form of dealing with the atrocities of breast cancer by means of MTT, Apoptotic, Western Blot Assays besides Transwell & Wound healing Assay. Reduction in the solid tumour in-vivo & near normalcy restoration of tissue section architecture from the BSO treated tumour sets are indicative of the better anti-tumorigenic potential of BSO.

**Methods:** BSO’s Solvent extraction was performed followed by its characterization. MTT aided cytotoxicity study of BSO alongside major components in PBMC & cancer cells while its efficacy was presented by flow cytometric ROS analysis, cell cycle arrest & apoptosis assessment. Anti-migratory potential evaluated by Wound Healing, Transwell Migration & Western Blot while the expression study of a wide range of proteins, miRNAs & the in-vivo studies undertaken climaxed the confirmation of the anti-cancerous potential of BSO.

**Results:** Comparatively reduced concentrations of TQ, TA & Carvacrol in BSO played a synergistic role to enhance apoptotic potential via Caspase 7 & 9, through enhanced ROS & expression of apoptotic family of proteins, miRNAs besides uplifting the anti-migratory perspective by effectively enhancing E-cad & downregulating lamellipodia, filopodia assembly & MMPs in MCF-7 & MDA-MB-231. Similar observations in-vivo outlined the therapeutic potential of BSO.

**Conclusion:** This study culminates isolation & processing of BSO in a simplified procedure, thereby aiming at a more lucrative paradigm to be accepted in contemporary phytomedicine research.

## INTRODUCTION

Accounting for 21.5% of all cancers, breast cancer holds its place as one of the most conventional cancer, affecting women globally so much so that it tops the chart for maximum deaths worldwide (Sharma et al., 2010). As the conventional therapies – chemo, multi-drug applications trigger colossal side effects in majority of patients, a quest to find a relatively safer & non-toxic therapy, traditional or folk medicines are being recognized parallel to modern therapeutics (Nurgali et al., 2018).

Originated in South eastern Asia, *Nigella sativa* has been used for medicinal purposes since centuries. Its seeds, known by several names in different cultures worldwide (Yimer et al., 2019b)– black cumin, kalojeera, kalonji etc and its oil have a long usage history in various medical systems and food (Yimer et al., 2019a). Although the seeds & oil extracted from *Nigella sativa* holds treasures of health benefits but not much study has been directed to explore its wholesome potential instead of testifying its individual components against deadly diseases like Cancer.

The present study took into consideration the solvent extraction of Black seed oil (BSO) & its consequent effect on the aggressive Triple negative breast cancer cell line MDA-MB 231 & HER2 negative MCF-7 which due to their non-responsive nature towards the conventional therapy became yet more difficult forms of breast cancer models to be dealt with (Holliday and Speirs, 2011). Several studies in this direction have so far failed to give a concluding backdrop to its role, thereby catering enormous attention. Though, the individual components of black seed oil-TQ, TA & Carvacrol (Luo et al., 2016; Shahbazian et al., 2015) are found to deliver anti-cancer properties, however their singular doses are much higher and expensive. Interestingly, our study highlights that when present in combination in the BSO even in minute proportions, these molecules cumulatively endeavour a strong anti-tumorigenic effect. In the current study, BSO has contributed to a significant rise in p53 expression in the MCF-7 cells which has further triggered the activation of caspases 7 and 9. On the other hand, BSO induced apoptosis in MDAMB-231 cells which have been observed to occur in a p53 independent fashion. Furthermore, a declining Bcl-2 expression and a consistent DNA fragmentation along with enhancement of Bax & PTEN in both the cell lines allowed the irreversible apoptotic undertaking alongside involvement of the caspases. Additionally, our findings also highlight the anti-migratory role of BSO in the breast cancer cells. Furthermore, our *in vitro* findings have also been validated *in vivo* which strongly justifies the anti-tumorigenic potential of BSO. An array of work has been done to exhibit the potential of black seed &amp; the oil extracted in curbing serious ailments. Several reports confirmed the metabolites of *N. sativa* seeds having therapeutic usage on cardiovascular, respiratory, immune, and endocrine systems (Gilani et al., 2004, El-Tahir and Bakeet, 2006, Ait Mbarek et al., 2007) Okasha et al., (2017) (Ait Mbarek et al., 2007; Gharby et al., 2015) reported the successful application of black seed oil as an adjuvant topical therapy, curing psoriasis. Moreover, several reports surfaced in the last few years, depicting the therapeutic potential of different integrant of black seeds. Bhattacharya et al., (2015) reported retardation of migrating breast cancer cells under the influence of Nano encapsulated Thymoquinone (Bhattacharya et al., 2015). Besides, Carvacrol was also reported to have shown therapeutic potential, alleviating proliferation & inducing apoptosis in two human colon cancer cell lines, HCT116 and LoVo (Fan et al., 2015).

Although much of the previously designed work has concentrated on the effect of *Nigella sativa* or its individual component, majorly Thymoquinone combating several disorders (Woo et al., 2012) or being used as an adjuvant to ease the same, yet there has not been any detailed & conclusive report on the molecular mechanism of orchestrated use of this phytochemical against breast cancer. Thus the present study is focused to point out the apoptotic & anti-migratory potential of BSO extract, which if harnessed in its absolute capacity can trigger a safer, & a greater bioavailable form of treatment against breast cancer cell lines - MDA-MB 231, MCF-7 without causing any toxicity to the normal cells in vitro & in-vivo, free from the usual setbacks of drug toxicity accompanying the contemporary medical therapeutic approaches employing individual integrants, thereby depriving the rest of the constituents forming the whole compound from exhibiting their phenomenal calibre.

## MATERIALS & METHODS

### Sample Collection & Isolation of essential oil from Black Cumin seed

Black Cumin seeds (or locally known as Kalo jeera) were collected directly from the local market. The seeds were washed with water and air dried. Then they were warmed in methanol at 50 °C, dried in vacuum and mechanically grinded by mortar-pestle and homogenised in laboratory homogenizer machine. The homogenised seeds were kept in water (10gm/500mL) as suspension and heated at 90°C for 6h. Then, after stirring at room temperature overnight at dark, the straw yellow solution was filtered out in vacuum filtration by Whitman filter paper. The solution was then extracted with n-hexane, diethyl ether and dichloromethane. The combined organic phase was dried over sodium sulphate and evaporated in vacuum. Finally straw coloured thick oil was obtained which was kept at 4°C for further use.

### Pharmacognostical analysis of Black Cumin seed

The pharmacognostic study is particularly important in authenticating medicinal plants so as to avoid any kind of adulteration & substitution of the essence of any kind of plant samples. The whole experimentation in this regard was done in collaboration with the Department of Pharmacognosy, Central Ayurveda Research Institute for Drug Development (CARIDD), Kolkata to lay down the standardization criteria perfectly. 200g of black cumin seeds were used for the said investigation.

### Characterization

- **High Resolution - mass spectrometry (HRMS)**: - This was employed to analyse the presence of Thymoquinone and other anti-cancerous components of black seed such as Carvacrol and Trans-Anethole attendant in the oil sample (Stock, 2017).
- **Gas chromatography-mass spectrometry (GCMS)**: - After the identification of important anti-cancerous components in the oil sample by HRMS, GC-MS experimentations were performed to ascertain these identifications and also to identify the components at a molecular level (Casuga et al., 2016). The standards used here were Thymoquinone, Trans-anethole & Carvacrol (all the 3 purchased from SIGMA-ALDRICH).

### Other Reagents

Details enumerated in Supplementary section S1.

### Cell line & Cell culture

Human breast cancer cell lines MCF-7 & MDA-MB 231 were purchased from National Centre for Cell Science (Pune, India). Human whole blood was collected from adult healthy volunteers with prior consent in heparinized vacutainer blood collection tubes (BD, Franklin Lakes, NJ) in accordance with Ahir et al., (2015) (Ahir et al., 2015). Details enumerated in Supplementary section S1.

### Cytotoxicity assay

Determination of the toxic potential of Black Seed Oil (BSO) was done with the aid of MTT (Sigma) to find out the percentage cell survival in case of MCF-7 & MDA-MB 231 cells on treatment with BSO. The assay was performed according to Upadhyay et al., (2019) (Upadhyay et al., 2019). Details enumerated in Supplementary section S1.

### Deciphering Cellular Apoptosis by Flow Cytometry

In order to determine the apoptotic dose of BSO on MCF-7 &MDA-MB 231 cells, apoptosis assay was performed in accordance to Ahir et al., (2015) (Ahir et al., 2015). Details enumerated in Supplementary section S1.

### Measurement of intracellular ROS

Sets of 2 * 10^6^ cells (approximately) were treated by BSO at a specific dosage for 24 h. This experimentation was carried out in accordance to Upadhyay et al., (2019) (Upadhyay et al., 2019). Details outlined in the Supplementary section S1.

### Cell Cycle Analysis by Flow Cytometer

Cell cycle phase distribution of nuclear DNA was determined using a 488 nm argon laser light source and 640 nm band pass filter (linear scale) equipped fluorescence detector on flow cytometer (Accuri C6 Plus Flow Cytometer, Beckton Dickinson,San Jose) and assisted by Accuri C6 plus software (BectoneDickinson, San Jose). Ten thousand total events were acquired and histogram display of DNA content (x-axis, PI fluorescence) versus counts (y-axis) was displayed.

### Cellular morphology examination by Scanning Electron Microscopy (SEM)

The morphology of breast carcinoma cells under different conditions were observed under Scanning Electron Microscope (SEM) by following Bhattacharya *et al*., 2015 (Bhattacharya et al., 2015). Details enumerated in Supplementary section S1.

### Wound healing assay

Cancer cell migration under different conditions was examined by bidirectional wound healing assay following the protocol enumerated by Upadhyay et al., (2019) (Upadhyay et al., 2019). Details outlined in Supplementary section S1.

### Transwell migration assay

Transwell migration assay was performed using cell culture inserts (BD Biosciences, Sparks, USA) with pores (8 mm) as stated by Upadhyay et al., (2019) (Upadhyay et al., 2019). Details outlined in the Supplementary section S1.

### Western blot analysis

Treated MCF-7 & MDA-MB-231 cells were analysed by means of western blotting in accordance to Upadhyay et al., (2019) (Upadhyay et al., 2019) & were incubated with primary antibodies - anti- β actin, p53, p65 (Nucleus), pERK, Cl Caspase 3, Cl Caspase 7, Cl Caspase 9, Cl PARP, PTEN, BAX, Bcl2, E-Cadherin, MMP-2 & MMP-9 (purchased from Santa Cruz Biotechnologies, USA). Details outlined in the Supplementary section S1.

### RNA isolation and quantitative real-time PCR

Total RNA isolation was carried out by Phenol Chloroform method utilizing TRIZOL Reagent and evaluation of the different RNA expression incorporated in this study was done following Upadhyay et al., (2019) (Upadhyay et al., 2019). Details in Supplementary section S1.

### Tumor regression in BALB/c model

8 BALB/c mice (8weeks old and weighing ~30 g) purchased from registered animal supplier (Regd No.:1443/PO/b/11/ CPCSEA) were housed in the animal house of Department of Physiology, University of Calcutta, Kolkata, and were maintained according to the guidelines of the Institutional Animal Ethical Committee (IAEC). The animals were kept in polypropylene cages and under controlled conditions of temperature and humidity with 12 h light/day cycles. They were given standard pellet diet, composed of 21% protein, 5% lipids, 4% crude fiber, 8% ash, 1% calcium, 0.6% phosphorus, 3.4% glucose, 2% vitamin, and 55% nitrogen-free extract (carbohydrates), purchased from Agro Corporation Pvt. Ltd., India and water ad libitum. After acclimatisation under the laboratory condition, they were randomly divided into 3 groups, each group having four mice (i) Untreated control group. 4T1tumor bearing set (which were injected with 4T1 cells in mammary fat pads), (ii) BSO treated set (T1) & (iii) BSO treated set (T2). Detailed experiments were performed using BSO at a dosage of T1 which represents 20 mg/kg while T2 that represents 30 mg/kg body weight (via oral gavaging). Tumor volume was measured every time dose was given and tumor weight evaluated following sacrifice of the mice & at the end of the experimental period of 15 days, animals were sacrificed by cervical dislocation. After sacrifice, the blood, serum and major organs (liver, kidney, spleen, lungs, heart and thymus) of the mice in the treatment and control groups were collected for further testing of toxicity by the measurement different toxicity markers.

### Serum toxicity assay

Details enumerated in the Supplementary section S1.

### Histopathological studies

This study was conducted in accordance to Upadhyay et al., (2019) (Upadhyay et al., 2019). Details enumerated in Supplementary section S1.

### Statistical investigation

The experimentations in general were performed in triplicates at least. The data were presented as the mean ± SD. The statistical significance was studied by one-way analysis of variance (ANOVA). The differences were considered to be statistically significant when the P values were less than 0.05 (P < 0.05).

## RESULTS

### Pharmacognostical Analysis of Black Cumin Seed, validating sample authentication

The Pharmacognostical analysis of Black Cumin seeds was carried out in the Department of Pharmacognosy, Central Ayurveda Research Institute for Drug Development, Kolkata. A total of 200g unprocessed, raw black seeds were used in the process. The entire set of investigations done in this area was in accordance to the Ayurvedic pharmacopoeia of India (API) Part I, Vol.I, 2001, Page:119 (Joshi et al., 2017) & the major focus of the study was the “seed” of Upakunchika, the Ayurvedic name of Black cumin seed (*Nigella sativa*) belonging to the family of Rananculaceae. The organoleptic characters of the seeds were observed to be Black (colour), Characteristic, Aromatic (odour) & Bitter (taste). The next set of identifications were done on the basis of Macroscopy [Fig.1(A)& (B)] & Powder Microscopy [Fig.1(C-K) wherein the former revealed that the seeds were small, flattened, oblong, angular, rugulose tubercular, pear shaped, one side was flat and the other convex. 0.3-0.4 cm long and 0.1-0.2 cm wide while the powder microscopy revealed that the seed powder exhibited epidermal cells Fig.1(C) consisting of elliptical, thick walled cells, brownish black parenchymatous cells containing oil globules Fig. 1(D, E), endosperm with oil globules Fig. 1(F), numerous oil globules Fig. 1(G), brownish content Fig. 1(H), aleurone grains, long fiber with narrow lumen Fig.1(I), prismatic calcium oxalate crystals Fig.1(J)& scalariform vessels Fig.1(K).

**Figure 1.**
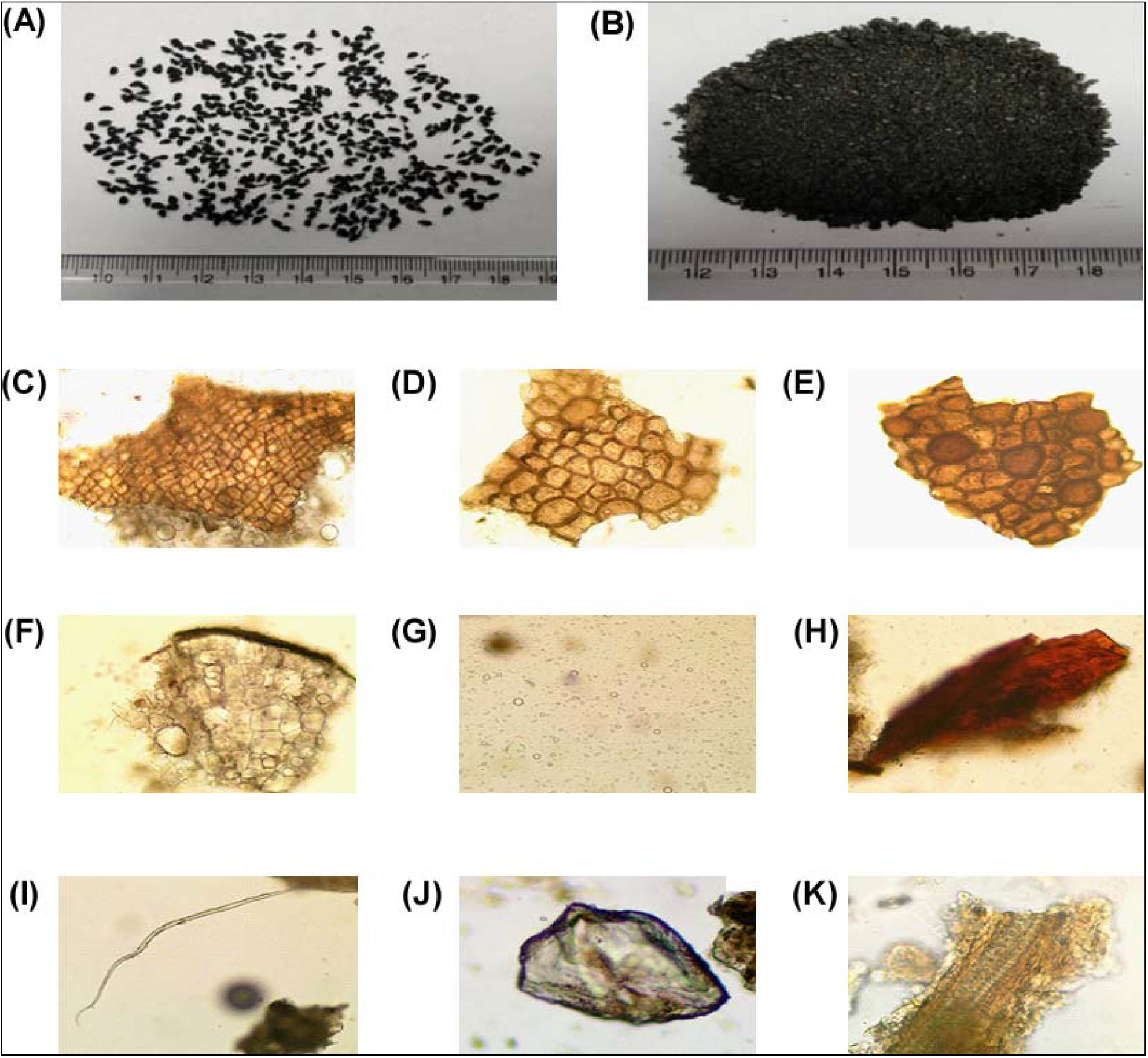
**(A) & (B)** Pharmacognostical Analysis of Black Cumin Seed: Macroscopy of Black Cumin Seeds. **(C)-(K)** Powder microscopy of Black Cumin seed: **(C)** Epidermis; **(D)&(E)** Parenchymatous cells; **(F)** Endosperm; **(G)** Oil globules; **(H)** Brownish content; **(I)** Long Fiber; **(J)** Calcium oxalate crystals; **(K)** Scalariform vessel

### Black seed Oil (BSO) Extraction & Characterization-Employing HRMS & GC-MS

In the present study, we have reported a very simple and effective process for isolation of essential oil from naturally occurring black seed. The schematic diagram showed in Fig. 2 described the isolation protocol of Black Seed Oil or BSO from the Cumin seeds.

**Figure 2.**
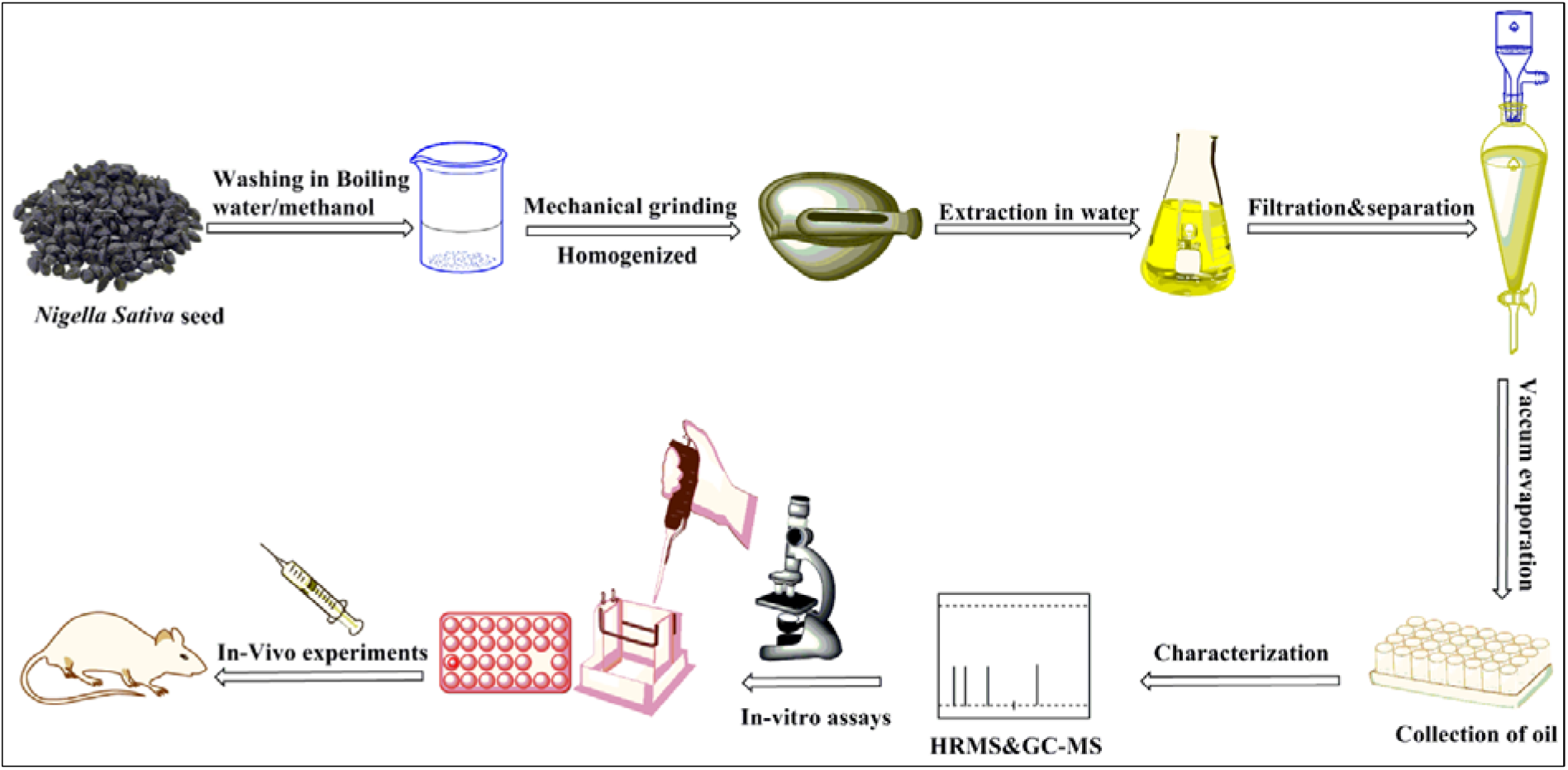
Schematic Representation of the Black Seed Oil (BSO) Extraction: BSO was extracted from Black Cumin seeds (Kala Jeera) & analysed using GC-MS & HRMS following which the oil was investigated to have any anti-cancerous properties *in-vitro* & *in-vivo*.

The oil was characterized by High-resolution mass spectra on a Xevo G2S/Q-Toff microTM spectrometer. Fig 3 showed HRMS of thymoquinone, Carvacrol and trans-anethole, vouching for a higher existence of thymoquinone in amounts more than either Carvacrol or trans-anethole. Most of these compounds were reported earlier as the primary compounds present in black seed. The extracted oil was directly used to observe its effect on the breast cancer cell lines as well as on animal models. The next characterization protocol in line was Gas chromatography–mass spectrometry (GC-MS) [Trace GC Ultra-Polaris Q] from Thermo Scientific. The pie chart highlights the compounds identified in the organic isolates from black seed extract [Fig. 4 (Lower Panel)]. Among them, the following were found in higher amounts: Thymoquinone (TQ), Carvacrol and Trans-Anethole (TA). Table 1 highlights the phytochemical fractions obtained from the GC-MS analysis of BSO.

**Figure 3.**
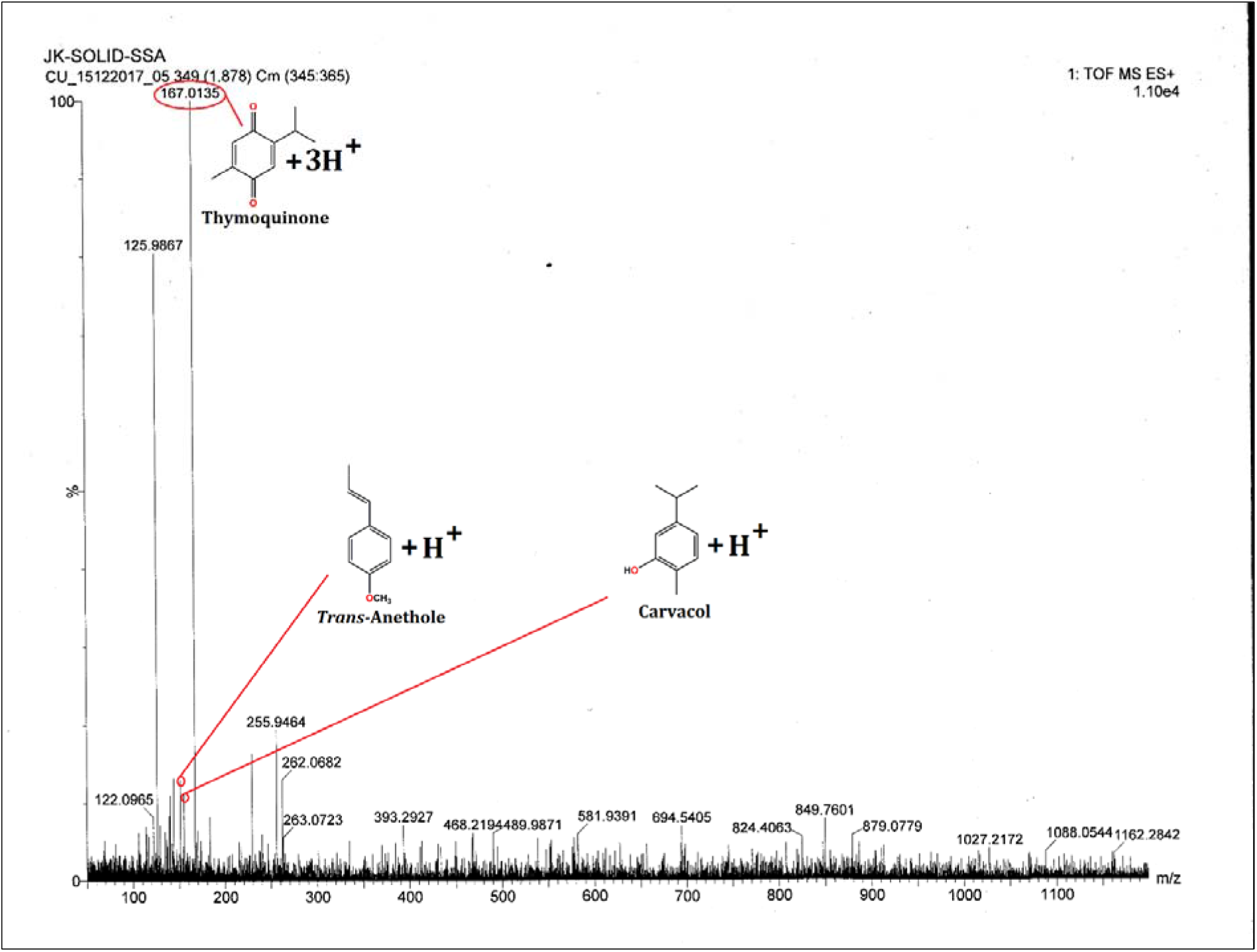
HRMS Data Representation: HRMS data showing the significant existence of Thymoquinone, Trans-Anethole & Carvacrol, contributing to the anti-migratory role of black seed extract.

**Figure 4.**
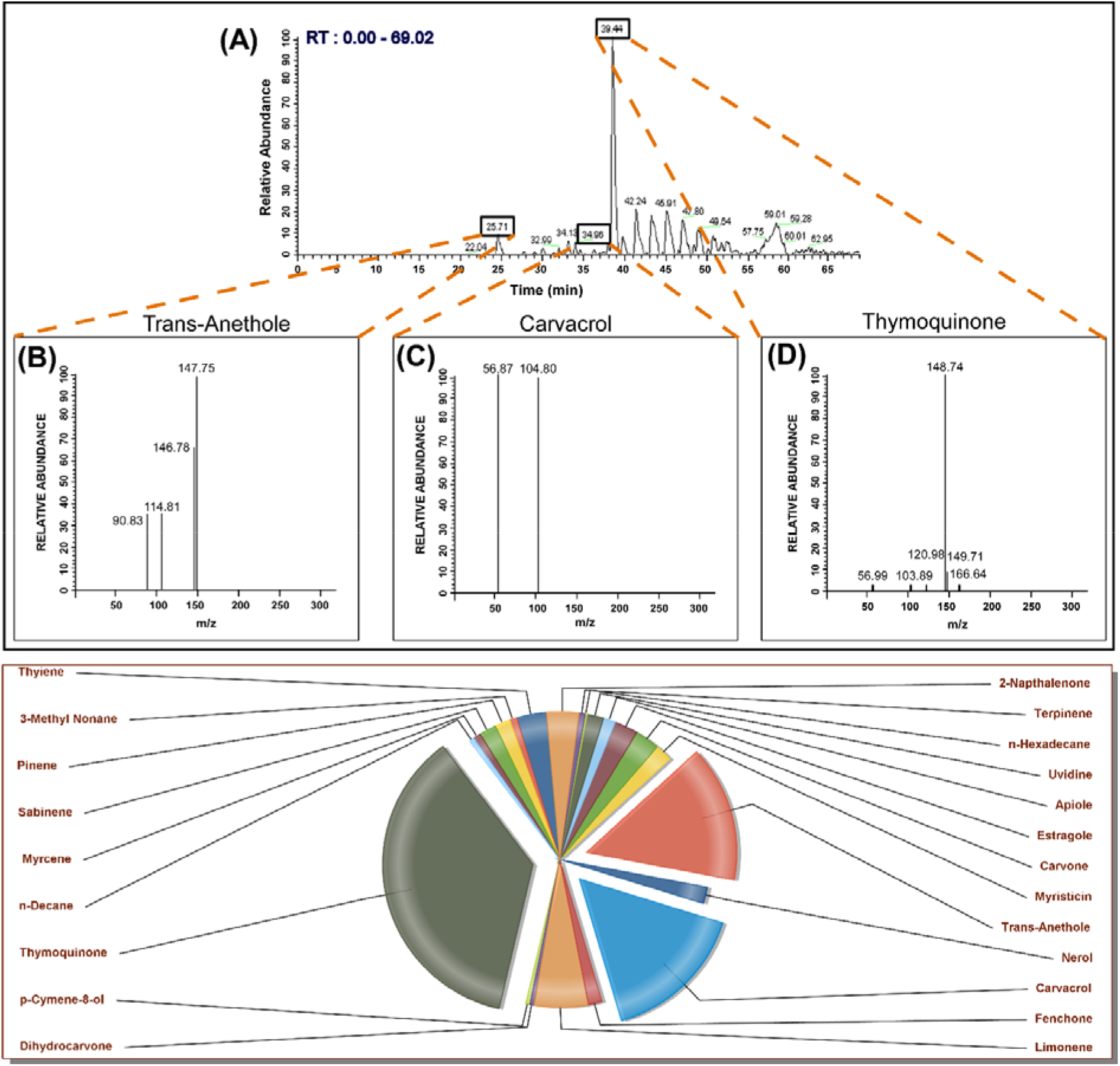
Interpretation of the various constituents of Black Seed Extract by means of **GC**-MS Analysis:[Upper Panel (A-D)] GC-MS Chromatogram of Extracted Oil from Black Seed; (A) GC-MS Chromatogram of the wholesome oil; (B) GC-MS Chromatograms showing the presence of Trans-Anethole at a Retention Time (RT) of 25.71 minutes, **(C)** Carvacrol at a Retention Time (RT) of 34.96 minutes & (D) Thymoquinone at a Retention Time (RT) of 39.44 minutes in the extracted Oil sample. The identification of the components was done in accordance to the Retention Indices & Mass Spectra of the samples authenticated & documented by the NIST library. Chart representing the components of Black Seed Extract, isolated from the Cumin seeds [Lower Panel].

**Table 1.**
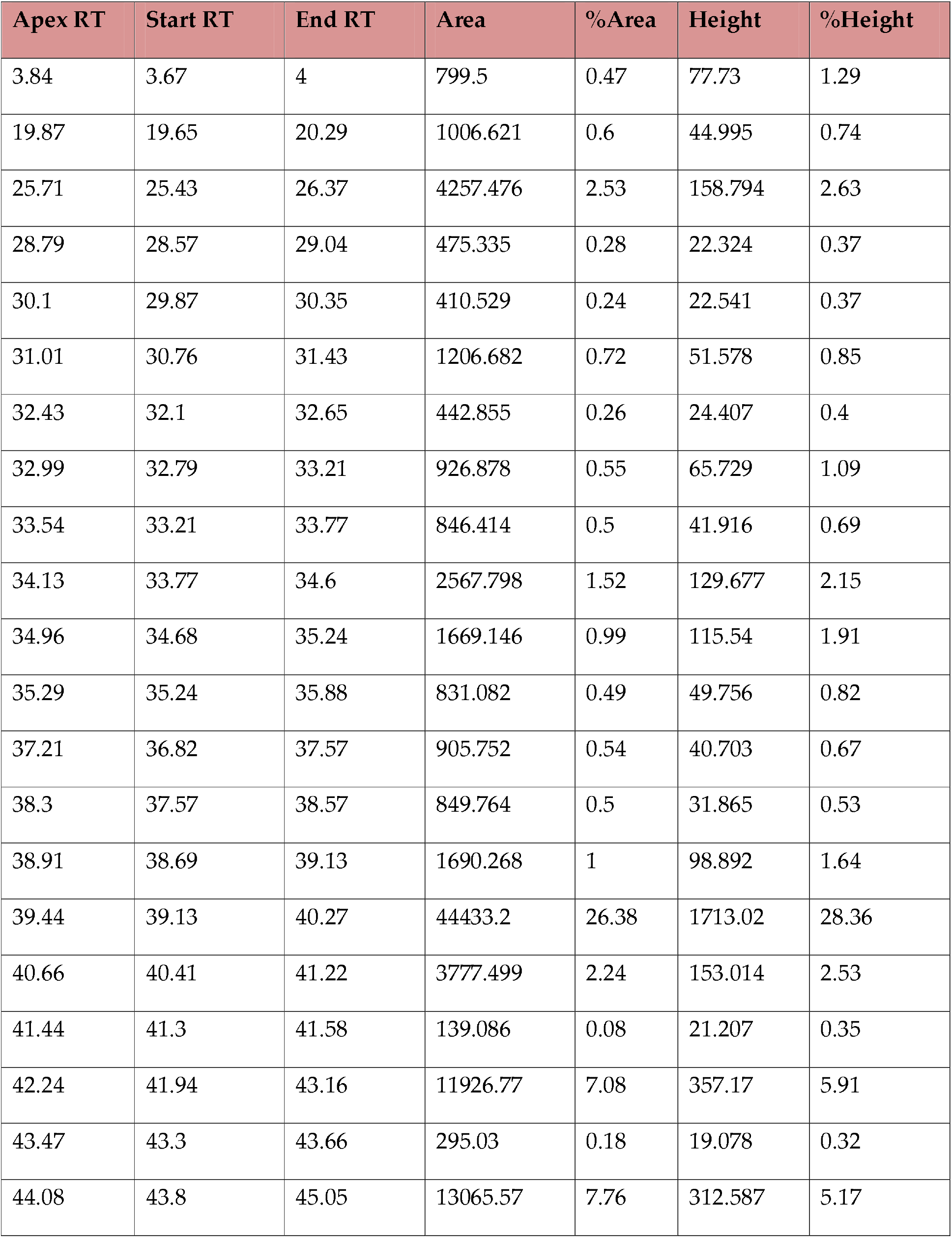
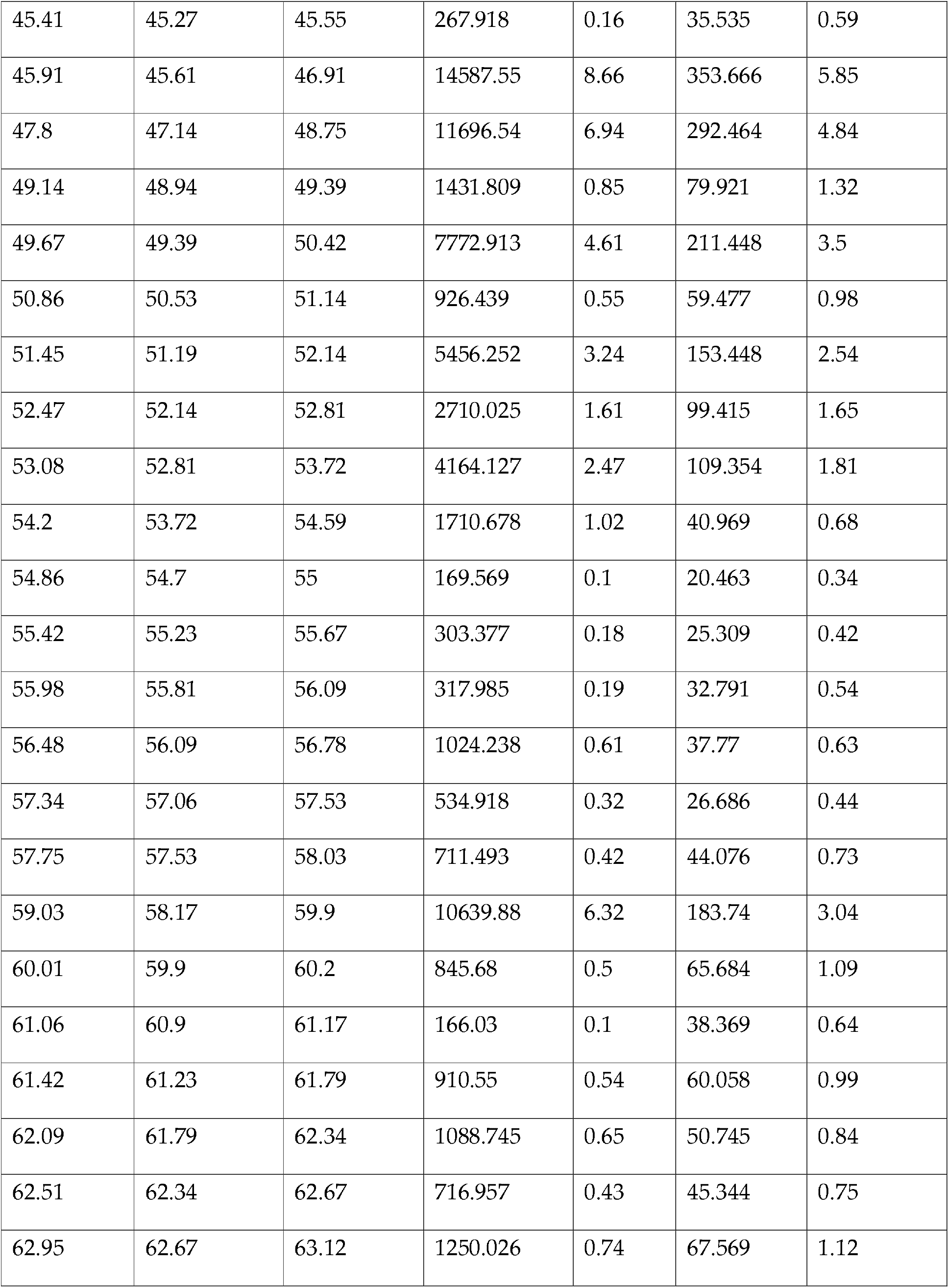
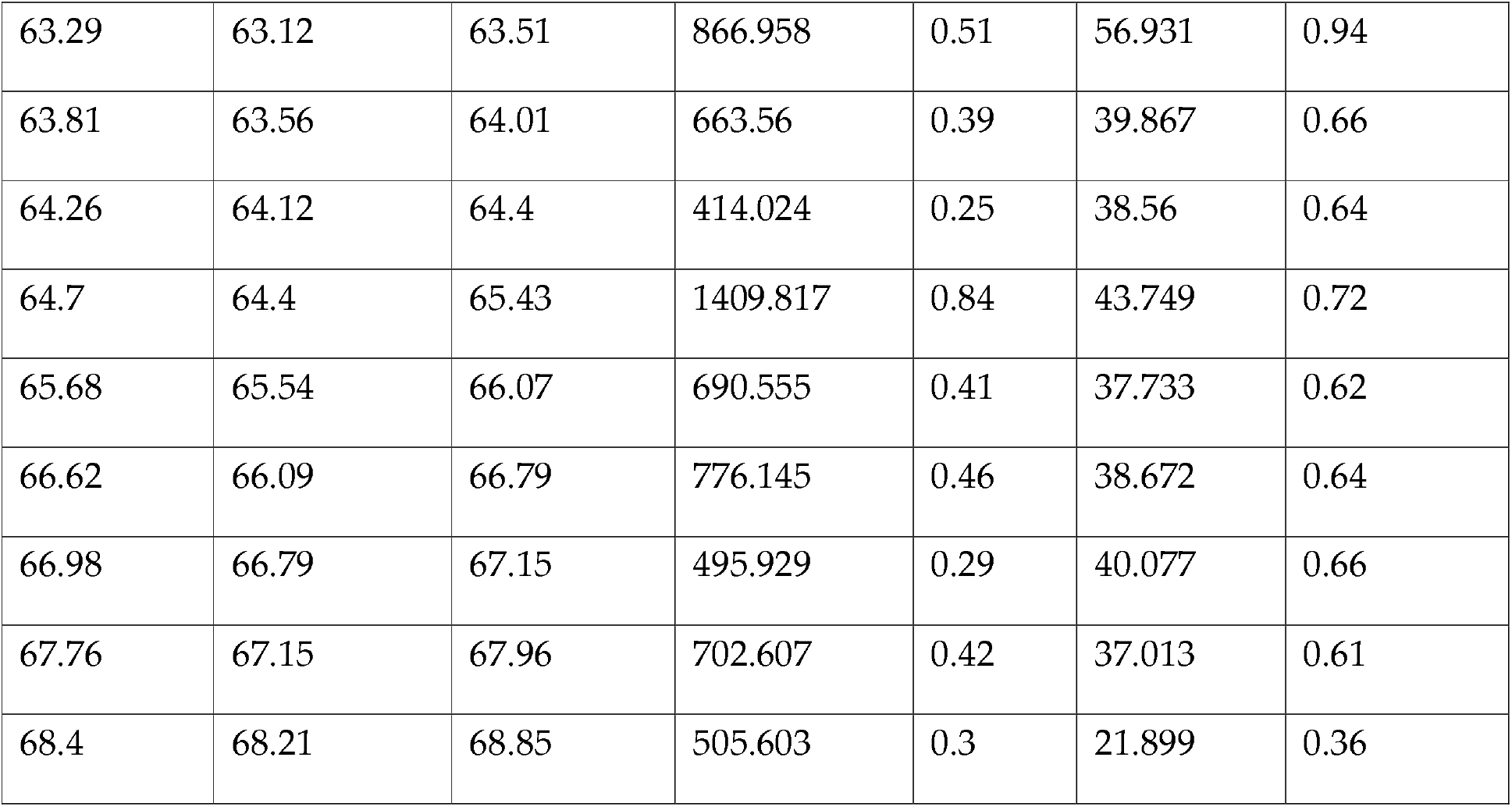
Characterization of Oil by GC-MS: List of Peaks obtained from the GC-MS Chromatogram (RT: 0.00 – 69.02; number of detected peaks: 53)

| Apex RT | Start RT | End RT | Area | %Area | Height | %Height |
| --- | --- | --- | --- | --- | --- | --- |
| 3.84 | 3.67 | 4 | 799.5 | 0.47 | 77.73 | 1.29 |
| 19.87 | 19.65 | 20.29 | 1006.621 | 0.6 | 44.995 | 0.74 |
| 25.71 | 25.43 | 26.37 | 4257.476 | 2.53 | 158.794 | 2.63 |
| 28.79 | 28.57 | 29.04 | 475.335 | 0.28 | 22.324 | 0.37 |
| 30.1 | 29.87 | 30.35 | 410.529 | 0.24 | 22.541 | 0.37 |
| 31.01 | 30.76 | 31.43 | 1206.682 | 0.72 | 51.578 | 0.85 |
| 32.43 | 32.1 | 32.65 | 442.855 | 0.26 | 24.407 | 0.4 |
| 32.99 | 32.79 | 33.21 | 926.878 | 0.55 | 65.729 | 1.09 |
| 33.54 | 33.21 | 33.77 | 846.414 | 0.5 | 41.916 | 0.69 |
| 34.13 | 33.77 | 34.6 | 2567.798 | 1.52 | 129.677 | 2.15 |
| 34.96 | 34.68 | 35.24 | 1669.146 | 0.99 | 115.54 | 1.91 |
| 35.29 | 35.24 | 35.88 | 831.082 | 0.49 | 49.756 | 0.82 |
| 37.21 | 36.82 | 37.57 | 905.752 | 0.54 | 40.703 | 0.67 |
| 38.3 | 37.57 | 38.57 | 849.764 | 0.5 | 31.865 | 0.53 |
| 38.91 | 38.69 | 39.13 | 1690.268 | 1 | 98.892 | 1.64 |
| 39.44 | 39.13 | 40.27 | 44433.2 | 26.38 | 1713.02 | 28.36 |
| 40.66 | 40.41 | 41.22 | 3777.499 | 2.24 | 153.014 | 2.53 |
| 41.44 | 41.3 | 41.58 | 139.086 | 0.08 | 21.207 | 0.35 |
| 42.24 | 41.94 | 43.16 | 11926.77 | 7.08 | 357.17 | 5.91 |
| 43.47 | 43.3 | 43.66 | 295.03 | 0.18 | 19.078 | 0.32 |
| 44.08 | 43.8 | 45.05 | 13065.57 | 7.76 | 312.587 | 5.17 |
| 45.41 | 45.27 | 45.55 | 267.918 | 0.16 | 35.535 | 0.59 |
| 45.91 | 45.61 | 46.91 | 14587.55 | 8.66 | 353.666 | 5.85 |
| 47.8 | 47.14 | 48.75 | 11696.54 | 6.94 | 292.464 | 4.84 |
| 49.14 | 48.94 | 49.39 | 1431.809 | 0.85 | 79.921 | 1.32 |
| 49.67 | 49.39 | 50.42 | 7772.913 | 4.61 | 211.448 | 3.5 |
| 50.86 | 50.53 | 51.14 | 926.439 | 0.55 | 59.477 | 0.98 |
| 51.45 | 51.19 | 52.14 | 5456.252 | 3.24 | 153.448 | 2.54 |
| 52.47 | 52.14 | 52.81 | 2710.025 | 1.61 | 99.415 | 1.65 |
| 53.08 | 52.81 | 53.72 | 4164.127 | 2.47 | 109.354 | 1.81 |
| 54.2 | 53.72 | 54.59 | 1710.678 | 1.02 | 40.969 | 0.68 |
| 54.86 | 54.7 | 55 | 169.569 | 0.1 | 20.463 | 0.34 |
| 55.42 | 55.23 | 55.67 | 303.377 | 0.18 | 25.309 | 0.42 |
| 55.98 | 55.81 | 56.09 | 317.985 | 0.19 | 32.791 | 0.54 |
| 56.48 | 56.09 | 56.78 | 1024.238 | 0.61 | 37.77 | 0.63 |
| 57.34 | 57.06 | 57.53 | 534.918 | 0.32 | 26.686 | 0.44 |
| 57.75 | 57.53 | 58.03 | 711.493 | 0.42 | 44.076 | 0.73 |
| 59.03 | 58.17 | 59.9 | 10639.88 | 6.32 | 183.74 | 3.04 |
| 60.01 | 59.9 | 60.2 | 845.68 | 0.5 | 65.684 | 1.09 |
| 61.06 | 60.9 | 61.17 | 166.03 | 0.1 | 38.369 | 0.64 |
| 61.42 | 61.23 | 61.79 | 910.55 | 0.54 | 60.058 | 0.99 |
| 62.09 | 61.79 | 62.34 | 1088.745 | 0.65 | 50.745 | 0.84 |
| 62.51 | 62.34 | 62.67 | 716.957 | 0.43 | 45.344 | 0.75 |
| 62.95 | 62.67 | 63.12 | 1250.026 | 0.74 | 67.569 | 1.12 |
| 63.29 | 63.12 | 63.51 | 866.958 | 0.51 | 56.931 | 0.94 |
| 63.81 | 63.56 | 64.01 | 663.56 | 0.39 | 39.867 | 0.66 |
| 64.26 | 64.12 | 64.4 | 414.024 | 0.25 | 38.56 | 0.64 |
| 64.7 | 64.4 | 65.43 | 1409.817 | 0.84 | 43.749 | 0.72 |
| 65.68 | 65.54 | 66.07 | 690.555 | 0.41 | 37.733 | 0.62 |
| 66.62 | 66.09 | 66.79 | 776.145 | 0.46 | 38.672 | 0.64 |
| 66.98 | 66.79 | 67.15 | 495.929 | 0.29 | 40.077 | 0.66 |
| 67.76 | 67.15 | 67.96 | 702.607 | 0.42 | 37.013 | 0.61 |
| 68.4 | 68.21 | 68.85 | 505.603 | 0.3 | 21.899 | 0.36 |

### BSO implemented Breast Cancer cytotoxicity in vitro while abrogating normal cell toxicity

Next, we evaluated the role of BSO & the three active ingredients individually on MCF-7, MDA-MB-231 as well as in Peripheral Blood Mononuclear Cells (PBMC), using MTT assay wherein we treated the cell lines with different concentrations of BSO, Thymoquinone (TQ), Trans-Anethole (TA) & Carvacrol. When exposed to the individual treatments, the cells outlined a substantially poor survival percentage as compared to BSO, conferring the credibility to the highly processed & purified components in contrast to BSO, housing a cocktail of the three mentioned ingredients in trace amounts, as highlighted by GC-MS [Fig. 4 (Lower Panel)] and including several other organic valuables in small proportion that works in consortium to achieve the desired cell death percentage. The concentrations at which the drugs were used ranged from 5-50 μg/mL & we observed a dose dependent cell death within 24 hours of BSO treatment in case of both the cell lines. Contrastingly, treatment with BSO also achieved a much higher cell survival percentage as compared to the other two cell lines in PBMC where the observed IC_50_of 50 μg/mL is sufficiently higher as compared to 20 μg/mL, 30 μg/mL & 35 μg/mL of TQ, TA & Carvacrol respectively [Fig 5. (A)]. Consequently, the Cytotoxicity profiling of the commercial integrant showed a considerably high percentage of cell death even at a very low dose in MDA-MB-231 with an IC_50_ of 10 μg/mL, 25 μg/mL, 20 μg/mL on treatment with TQ, TA & Carvacrol respectively as compared to an IC50 of 30 μg/mL BSO treatment [Fig 5. (B)]. A similar line of observation surfaced on treatment of MCF-7 cells with the three active ingredients & BSO wherein the IC50 values noted for BSO was 25 μg/mL, TQ was 5 μg/mL & both TA & Carvacrol were 25 μg/mL each [Fig 5. (C)]. the red arrow indicates the IC_50_values for each graph. After screening various dosages in this study, we chose a dose below the IC_50_values, i.e., 15 μg/mL of BSO to carry out further investigation on its anti-migratory potential in case of both MCF-7 &MDA-MB-231 while rest of the experimentations were performed with the IC_50_ doses only.

**Figure 5.**
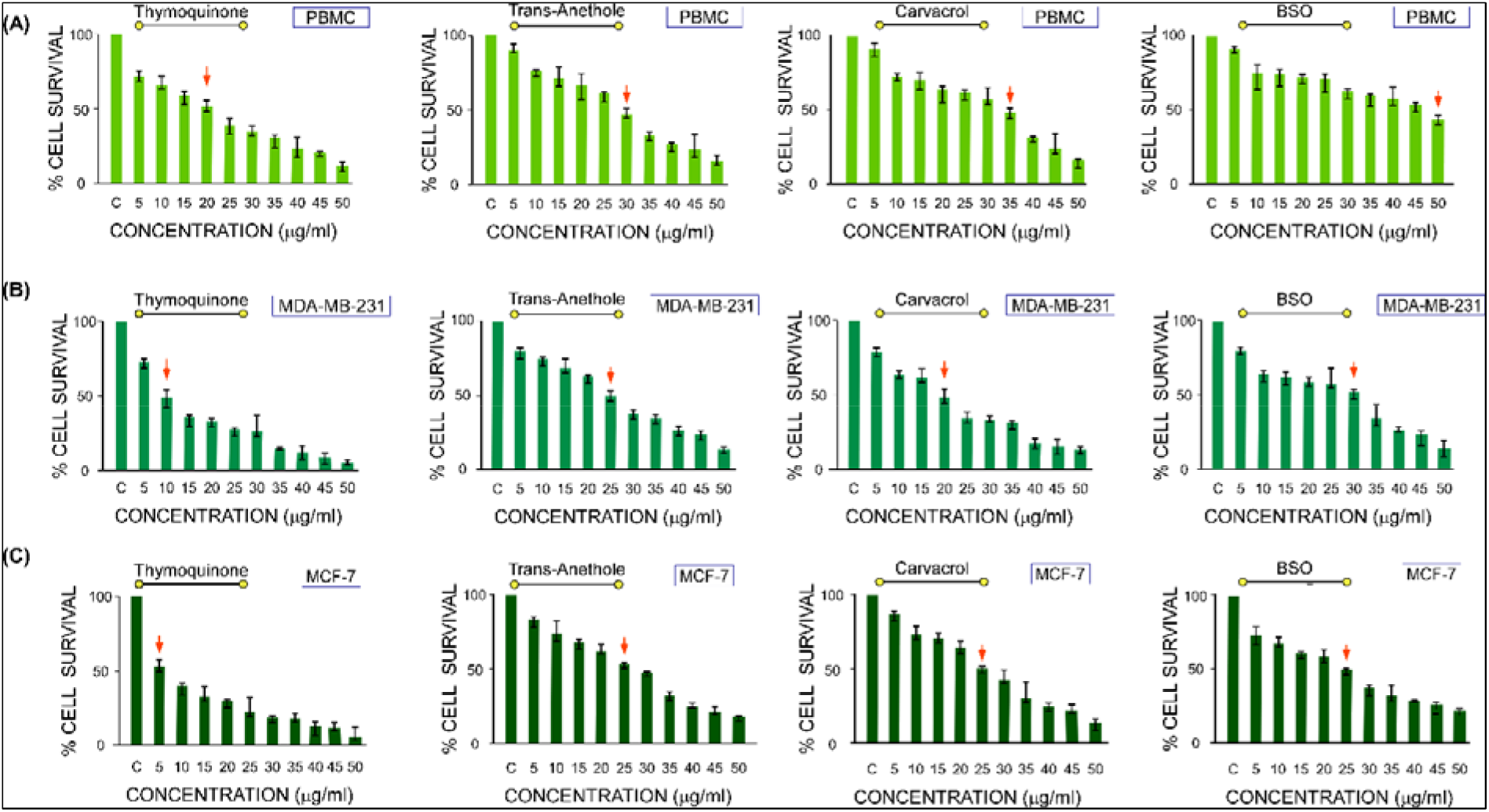
Cytotoxic effects of BSO & its chief integrant: **[A]** Graphical Representation, showing Percentage Cell Survival of Peripheral Blood Mononuclear Cells (PBMC); **(B)** MDA-MB-231 & (C) MCF-7 when treated with Thymoquinone (TQ), Trans-Anethole (TA), and Carvacrol individually & BSO in a dose dependant manner with the aid of MTT for 24 hours. The indicated dosages range from 5-50 μg/mL. The cell viability of Control Set was considered to be 100%. The red arrow indicates the IC50 values for each graph. All data shown are representative of at least 3 independent experiments.

### BSO induced ROS mediated apoptotic cell death in MDA-MB 231 & MCF-7 cells

Following the Cytotoxicity assay by MTT, we performed cell cycle analysis in control & BSO treated MCF-7 [Fig.6 (A).I] & MDA-MB-231 cells [Fig.6 (A). II] To check for any association between cell cycle arrest & cell demise as part of BSO treatment. As instantiated by Fig.6 (A).I & II, the 24 hour BSO treated cells showed a clear increment in the sub-G0/G1 peak which was evidently higher as compared to their untreated counterpart indicating towards significant cell death. Furthermore, the induction of significant G2/M phase cell cycle arrest was observed on BSO treatment.

**Figure 6.**
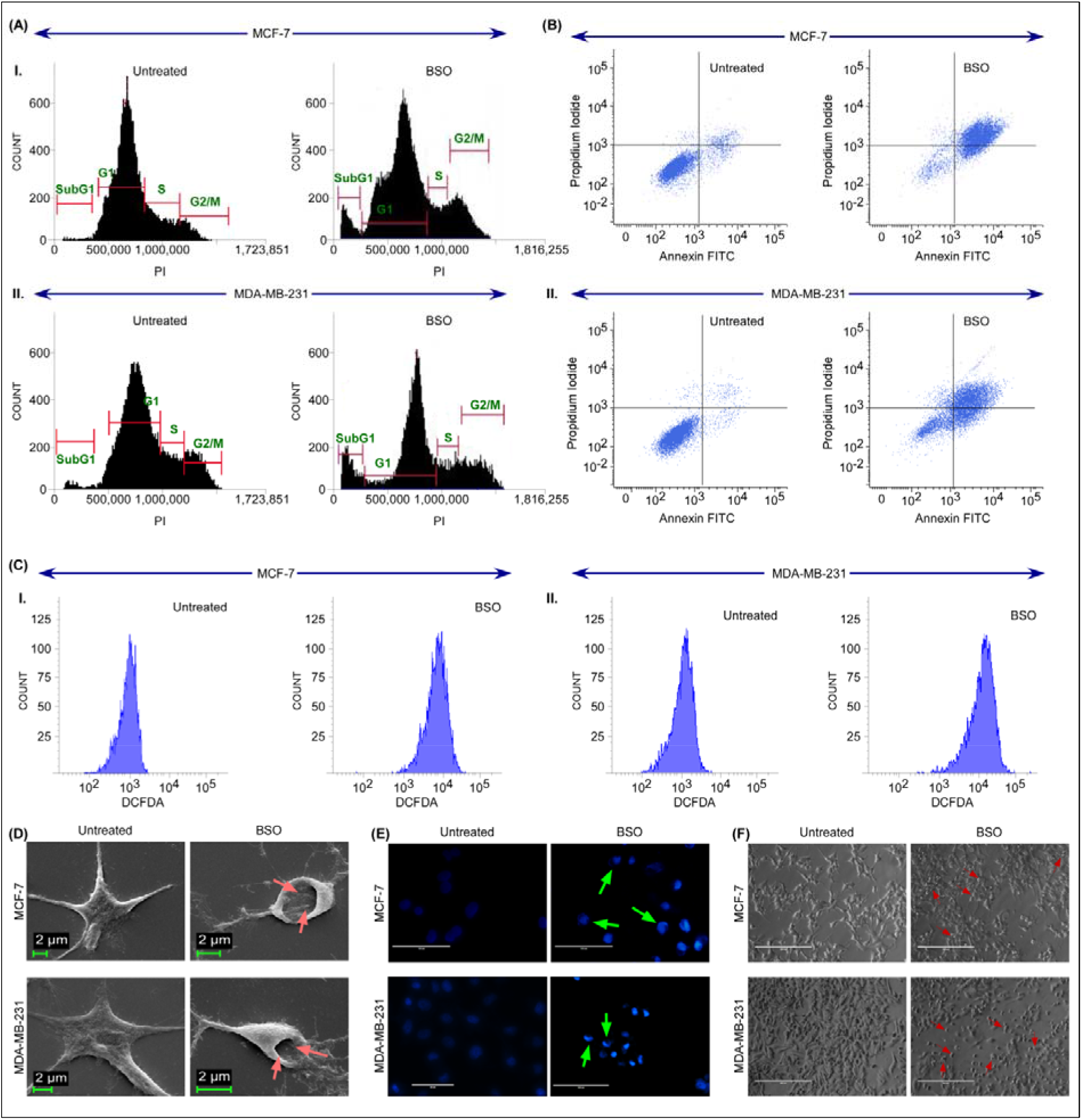
Induction of Apoptosis in Breast Cancer cells: Cell cycle phase distribution of nuclear DNA was determined by Flow Cytometric Assay in case of **(A) I.** MCF-7 & **(A) II.** MDA-MB-231 cells treated with 25 μg/mL BSO in case of MCF-7 & 30μg/mL for MDA-MB-231. Plots display the DNA content (PI fluorescence along the x-axis) versus cell count (along the y-axis). Induction of Apoptosis by BSO in **(B) I.** MCF-7 & **(B) II.** MDA-MB-231. Flow cytometric data measured by Annexin/PI showcases apoptotic and necrotic cell populations when treated with BSO for 24h. Dual parameter dot plot of FITC fluorescence (x-axis) versus PI fluorescence (y-axis) has been shown in logarithmic fluorescence to demarcate cells in different phase: (lower right quadrant) early apoptosis, (upper right quadrant) late apoptosis, and (upper left quadrant) necrosis cells. Flow Cytometric Analysis of intracellular ROS level by measuring the DCFDA-fluorescence intensity in **(C) I.** MCF-7 & **(C) II.** MDA-MB-231 for 12 hr in both untreated & treatment conditions i.e., 25 μg/mL BSO in MCF-7 & 30μg/mL for MDA-MB-231. **[(D) Right]** Cellular Morphology analysis by SEM micrographs of MCF-7 (upper) & MDA-MD-231 (lower) showing the cells in their late apoptosis phase, wherein the cells are visibly damaged due to apoptosis induction on BSO treatment. The arrows are indicative of the same. **[D) Left]** However, the untreated cells do not show any signs of such morphological discrepancies in both the cell lines. **[(E) Right]** Phase Contrast Images of MCF-7 (upper) & MDA-MD-231 (lower), indicating early apoptotic cells, suggestive of chromatin condensation & nuclear fragmentation (Green Arrows indicate the same) on staining with Hoechst 33342 (blue) **[(E) Left]** while no such observations were reported in case of untreated cells. **[(F) Right]** Phase contrast images, showing cell death by treatment with BSO in both breast cancer cell lines MCF-7 (upper) & MDA-MD-231 (lower). The same dosages are used as stated above. The red arrows pin-points the area of cell death. Values are mean ± SEM of three independent experiments in each case or representative of typical experiment. **P <0.01, and ***P <0.001.Each data is the mean ± SD (n=3); *P<0.01, **P<0.001, and ***P<0.0001.

In order to confirm the mode of cell death in case of the breast cancer cells under investigation, Annexin-V-PI assay was performed which highlighted almost 45% cell death on treatment with 25 μg/mL BSO in MCF-7 cells [Fig.6 (B).I]. The IC_50_ dosage of 30 μg/mL used in case of MDA-MB-231 garnered almost 46% cell death [Fig.6 (B).II] reinstating the role of BSO in inducing apoptosis. The observations revealed that apoptotic cell death also resulted in cell membrane blebbing & formation of apoptotic bodies as evident by the morphological analysis using Scanning Electron Microscope (SEM) [Fig.6 (D)] accompanied nuclear fragmentation [Fig.6 (E)] on treatment with BSO as well. Several studies revealed that elevated levels of ROS promote cytochrome c to be released into the cytoplasm and triggers programmed cell death (Vacca et al., 2006). In the present experimental set up, we observed that the treatment with BSO promoted ROS generation in both the cell lines. BSO was observed to be instrumental in elevating the level of ROS in MCF-7 [Fig.6 (C).Left]& MDA-MB-231 cells [Fig.6 (C).Left], thereby causing the inevitable cell demise. The changes in the morphology of the cells were investigated with the help of Hoechst 33342 (blue) which seconds the nuclear fragmentation creeping in the cell as a part of the apoptotic cell death regime [Fig.(6) E]. Besides, phase contrast microscopic images revealed a substantial difference in cell death in treated sets as compared to the untreated ones [Fig.(6) F]. The arrows are indicative of the morphological changes & cellular demise.

### BSO attenuated migratory potential in Breast Cancer cells

After ensuring the role of BSO as an apoptotic agent, we were next prompted to elucidate its role in cancer cell migration. The effect of BSO on breast cancer cell migration was examined by bidirectional wound healing assay wherein both the cell lines were treated with a dose of 15μg/mL BSO & the rate of migration observed 24 h after treatment on MCF-7 [Fig.7 (A) Right Panel]& MDA-MB-231 [Fig.7 (A) Left Panel] cell lines were noted. The reason behind selecting the particular dosage lies in its capability to result in sufficient cell death, thus proving to be an optimum dose for the experimentations being carried out here. These results marked the anti-migratory potential of BSO as depicted by Fig. 7 (A) which showed that in the BSO treated cell sets, the wound closure was hardly visible, thereby demonstrating the anti-migratory potential of BSO *in vitro*. Further, the mentioned findings have been supported by a graphical representation, exhibiting the percentage cell migration at 0 & 24 hours after treatment with 15 μg/mL BSO Fig. 7 (B). Next, the Transwell Assay was performed, where MCF-7 [Fig.7 (C) Left Panel] & MDA-MB-231 cell lines [Fig.7 (C) Right Panel] were exposed to 15 μg/mL BSO treatment respectively. A graphical representation further sketched the percentage of cell migration in both the cell lines Fig. 7 (D). SEM micrographs revealed the potential of BSO in attempting to limit the formation of Lamellipodia, the ribbon-like, flat cellular protrusions formed at the periphery of a moving cell& filopodia, the slender, actin-rich cell membrane extensions which enhances the cell motility, thereby curbing the two major stimulants of Cancer cell migration. Thus, these results attempted to establish BSO’s amplitude in serving a pivotal role as an anti-migratory formulation in breast cancer as in the case of MCF-7 [Fig.7 (E) Upper Panel]&MDA-MB-231[Fig.7 (E) Lower Panel] when treated with 15 μg/mL BSO. The migratory markers such as E-Cadherin, MMP-2 & MMP-9 to BSO treatment were also taken into consideration. Western blotting analysis represented the changes in MCF-7 [Fig.7 (F) Left Panel]& MDA-MB-231 [Fig.7 (F) Right Panel]. While E-Cadherin showed a considerable upregulation in the treated sets of both the cell lines, we observed downregulation in the expressions of MMP-2 & 9, thereby resonating their relationship with breast cancer prognosis.

**Figure 7.**
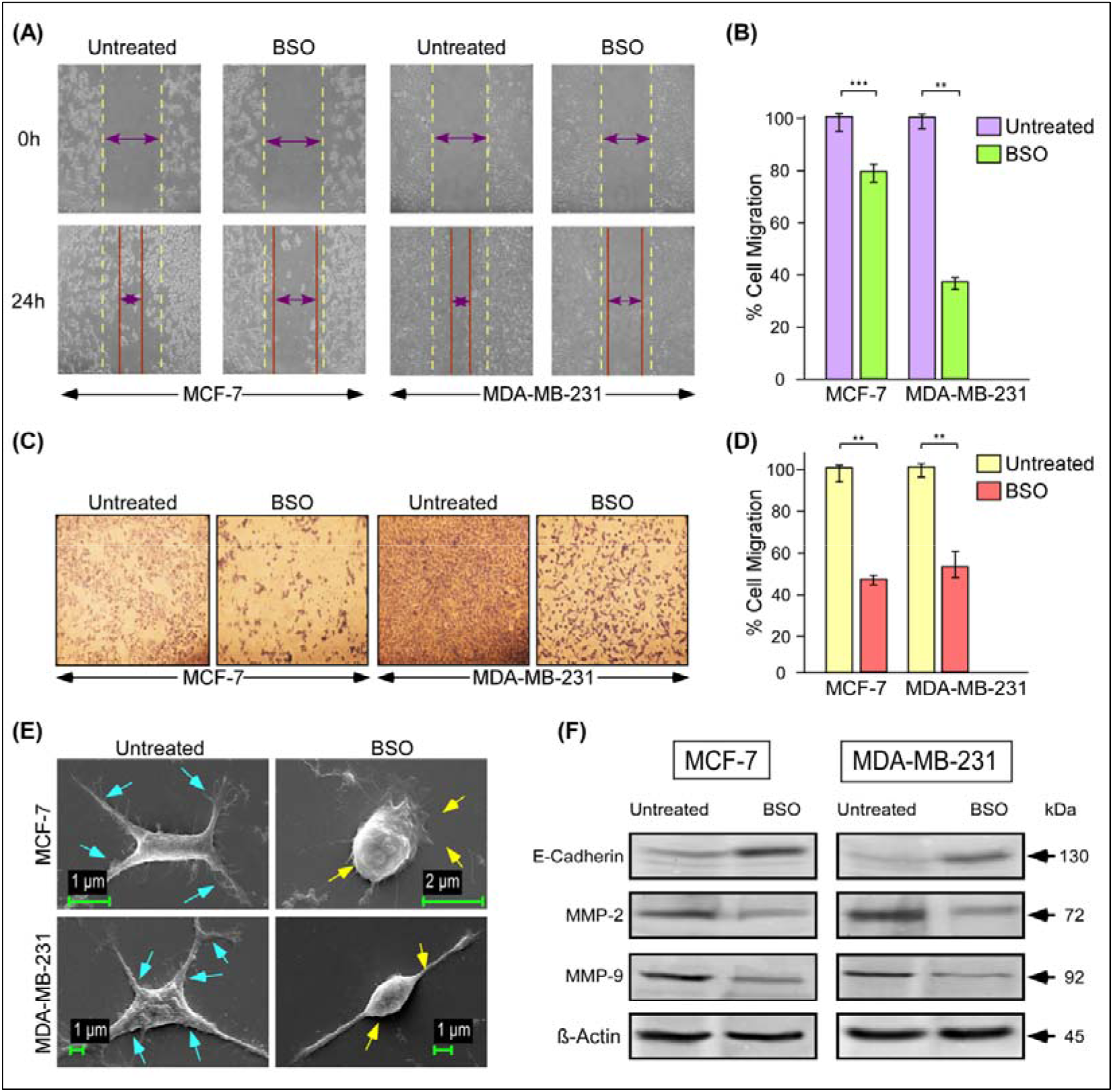
BSO curbs migration of MCF-7 &MDA-MB-231 cells *in vitro*: **(A)** Phase Contrast images, exhibiting the anti-migratory effect of BSO, evidently portrayed by the changing migratory patterns in the bi-directional wound healing assay in case of (left panel) MCF-7 & (right panel) MDA-MB-231. (B) Further, a graphical representation of the percentage cell migration at 0 & 24 hours showed the effect of BSO. (C) The percentage migration for Control cells is considered 100% in this experiment & the comparisons drawn accordingly. Phase contrast images showing the stained (Left Panel) MCF-7 cells & (Right Panel) MDA-MB-231 cells as obtained from the Transwell migration assay in control and treatment with BSO. (D) A graphical representation of the same shows the Percentage of cell migration of MCF-7 & MDA-MB-231 under the said conditions. **(E)** SEM micrographs of **(Left Panel)** MCF-7 & **(Right Panel)** MDA-MB-231 highlighting the changes in the Lamellipodia & Filopodia structures. The arrows pin point the structural changes. **(F)** Western Blot analysis to showcase the expression of E-Cadherin, MMP-2 & MMP-9 using β-actin as an internal control. Values are mean ± SEM of three independent experiments in each case or representative of typical experiment. **P <0.01, and ***P<0.0001.

### BSO mediated changes in an array of Protein & miRNA expressions thereby facilitating breast cancer cellular demise

Western Blot Analysis was performed to ascertain the changes in expression of Bax, Bcl-2, p53, PTEN, p65 (nuclear), Cleaved (Cl) Caspase 9 & 7 [Fig.8 (A)] in BSO treated MCF-7 cells. The dosage used here was 25 μg/mL. A prominent increase in the expression of Bax & p53 with an upgraded expression of PTEN was evident towards the induction of apoptosis. However, conversely, a decreased Bcl-2 expression in BSO treated MCF-7 highlights the Bax/Bcl-2 ratio that remained intact even in this case, allowing the cells to embrace apoptosis. Additionally, the loss of Bcl-2 family proteins gave us much insight into the reduced nuclear expression of p65 in this case. Furthermore, we probed into the expression changes taking place in Caspase 9 & 7 which showed a concrete upregulation in the BSO treated MCF-7 cells as compared to their untreated counterparts which in turn confirmed the apoptotic cascade. A bar graph obtained from the qRT-PCR analysis depicted the changes in RNA expression of Bax, Bcl-2, and p53 [Fig.8 (B)].

**Figure 8.**
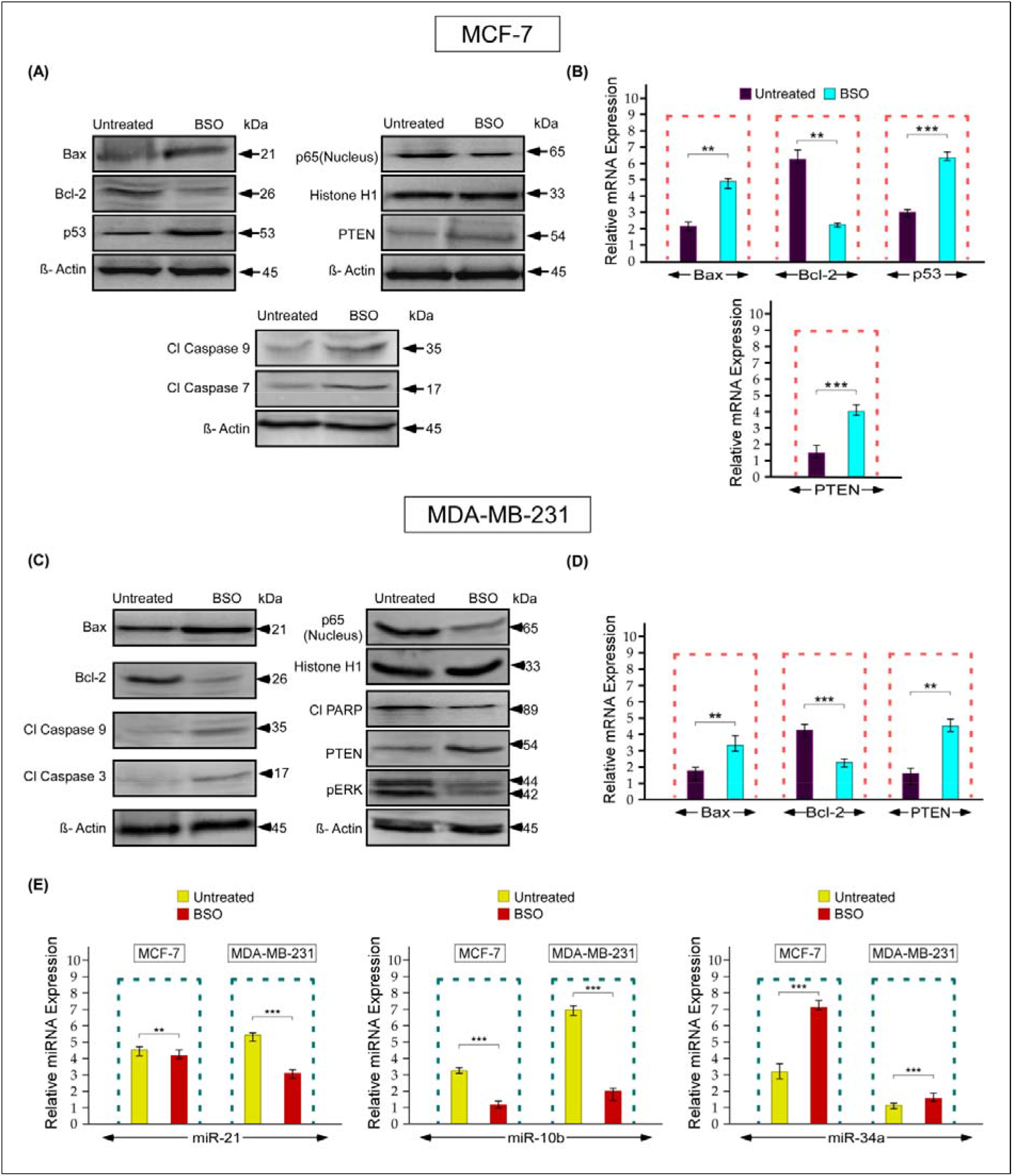
Triggering a varied change of expression in an array of proteins & miRNAs on an aftermath of BSO treatment: (A) Western Blot Analysis in MCF-7 cells showed transformations in the expression pattern of several proteins-Bax, Bcl-2, p53, PTEN, p65 (nuclear), Cleaved (Cl) Caspase 9 & Cleaved (Cl) Caspase 7 when treated with 15 μg/mL BSO. β-actin and Histone H1 was used as an internal control in this case. (B) Graphical representation showing the relative mRNA Expression of Bax, Bcl-2 and p53 on similar treatment in MCF-cells. 18s was used as the housekeeping gene in this set of qRT-PCR experiments. (C) The changes in the expression of Bax, Bcl-2, p65 (nuclear), PTEN, pERK, cl Caspase 9, cl Caspase 3 &Cleaved (Cl) PARP were evaluated by means of Western Blot Analysis on treatment of MDA-MB-231 cells with 15 μg/mL BSO. β-actin and Histone H1 served as the internal control in this case. (D) qRT-PCR results illustrated the mRNA Expression patterns of Bax, Bcl-2, and PTEN in the MDA-MB-231 cells on treatment with the above mentioned dosage of BSO. (E) Graphical representation of miRNA Expression pattern in case of (Left Panel) miR-21, (Middle Panel) miR-10b & (Right Panel) miR-34a by qRT-PCR on exposing both the breast cancer cells to a treatment of 15 μg/mL BSO. Values are mean ± SEM of three independent experiments in each case or representative of typical experiment. **P <0.01, and ***P <0.001.

From Fig.8 (C), we observed an enhanced Bax & PTEN expression contributing to apoptosis in MDAMB-231 cells but in a p53 independent manner. The reduced Bcl-2 & p65 expression (nuclear) again indicated that BSO treatment facilitated apoptotic cell death *in vitro*. Besides, a profound reduction in the pERK expression aided cell death by BSO given the affinity of pERK for contributing to the extreme invasive nature of breast cancer cell. Additionally, we observed a steady up-regulation of Cl caspase 9 & 3 in BSO treated MDA-MB-231 cells which directed the apoptotic death in the mentioned cell line. Also was evident the cleavage of PARP induced in the treated sets which culminated in programmed cell death. A bar graph was etched in support of the qRT-PCR analysis which showed the RNA expressions of Bax, Bcl-2 and PTEN [Fig.8 (D)]. This study also brought to light an important contribution of BSO in the form of its ability to modulate expressions of small non-coding RNA molecule also known as micro RNA or miRNA & the same was exhibited with the help of qRT-PCR analysis [Fig.8 (E)]. The expression of miR-21 was seen to be decreased in case of BSO treated cells in both the cell lines which have been previously studied to have been negatively regulating tumour suppressors (Shen et al., 2014). Also, another well documented oncomiR aiding cancer cell migration showed reduction in its expression upon BSO treatment-miR-10b. We also studied the expression of miR-34a & reported a steady upliftment in case of MCF-7 & a negligible rise in MDA-MB-231 pertaining to its contribution of negatively regulating the anti-apoptotic proteins, thereby assisting apoptosis.

### In-vivo validation of tumor restraining effect of BSO on oral administration & scrutinizing the histological changes in peripheral tissues and tumor sections

In order to ascertain that the treatment of BSO houses ability to curb growth of solid tumors *in vivo*, we used female BALB/c mice (Weight-30g) in which 4T1 mammary carcinoma cells were injected into the mammary fat pads and typically formed a solid tumor which became visible 1 week after the injection as shown in Fig. 9 (A) [‘No treatment’]. The tumour size kept on increasing till the time of sacrifice. BSO was then orally administered in the mice every alternate day till the next 20 days &[Fig. 9 (A) ‘T1’ & ‘T2’] documented the changes in the tumor size incurred after oral gavaging of BSO where T1= 20 mg/kg while T2= 30 mg/kg body weight of mice. As a consequence of treatment by BSO, there was a considerable reduction in the Tumor Volume [Fig. 9 (B)] & Weight [Fig. 9 (C)]. The following set of experimentations were performed using T2 i.e., 30 mg/kg body weight of mice. Furthermore, on performing qRT-PCR analysis of Bax & Bcl-2, we found an enhanced Bax expression in along with the reduction of Bcl-2 which confirmed the compromised state of anti-apoptotic proteins by virtue of BSO treatment [Fig. 9 (D)]. The anti-tumour efficacy of the BSO treatment was reinforced by the results of the study of H&E stained sections of tumor from Tumor and treatment groups [Fig. 9 (E)]. BSO treatment resulted in a decrease in the neoplastic foci within the tumor of BSO treated animals as compared to those that did not receive BSO. Further, atrophying cancer cells with necrotic zones with less focal haemorrhage was evident in BSO group as compared to only tumour-bearing group. The nuclear to cytoplasmic area also appeared to be lower in BSO treated tumours indicating a reduction in the proliferation capacity in tumors treated with BSO. Heart, spleen, lungs, liver and kidneys harvested from treated animals were processed into 5-7 μm sections and stained with haematoxylin and eosin [Fig. 9 (F)]. Microscopic examination revealed that tumour development induced a gross deterioration of hepatic architecture demarcated by degeneration of the central vein, atrophic cells and inflammatory infiltrates. The renal morphology was also disrupted in tumour bearing mice as was evident form the increased urinary space, and cellular atrophy. Further, the cyto-architecture of the cardiac tissue also showed degenerative changes in the tumour bearing animals. Interestingly, the lung tissue sections from tumour bearing showed increased bronchial atrophy, inflammatory infiltrate and also zones of hyper-cellularity which might be indicative of secondary tumour growth. Further, study of the splenic sections revealed areas of cellular atrophy and other indicators of loss of normal architecture in the tumour bearing mice. It was noteworthy that treatment with BSO appeared to reduce the degenerative morphological changes with the extent of architectural damage being apparently lower in the tissue sections of BSO treated mice.

**Figure 9.**
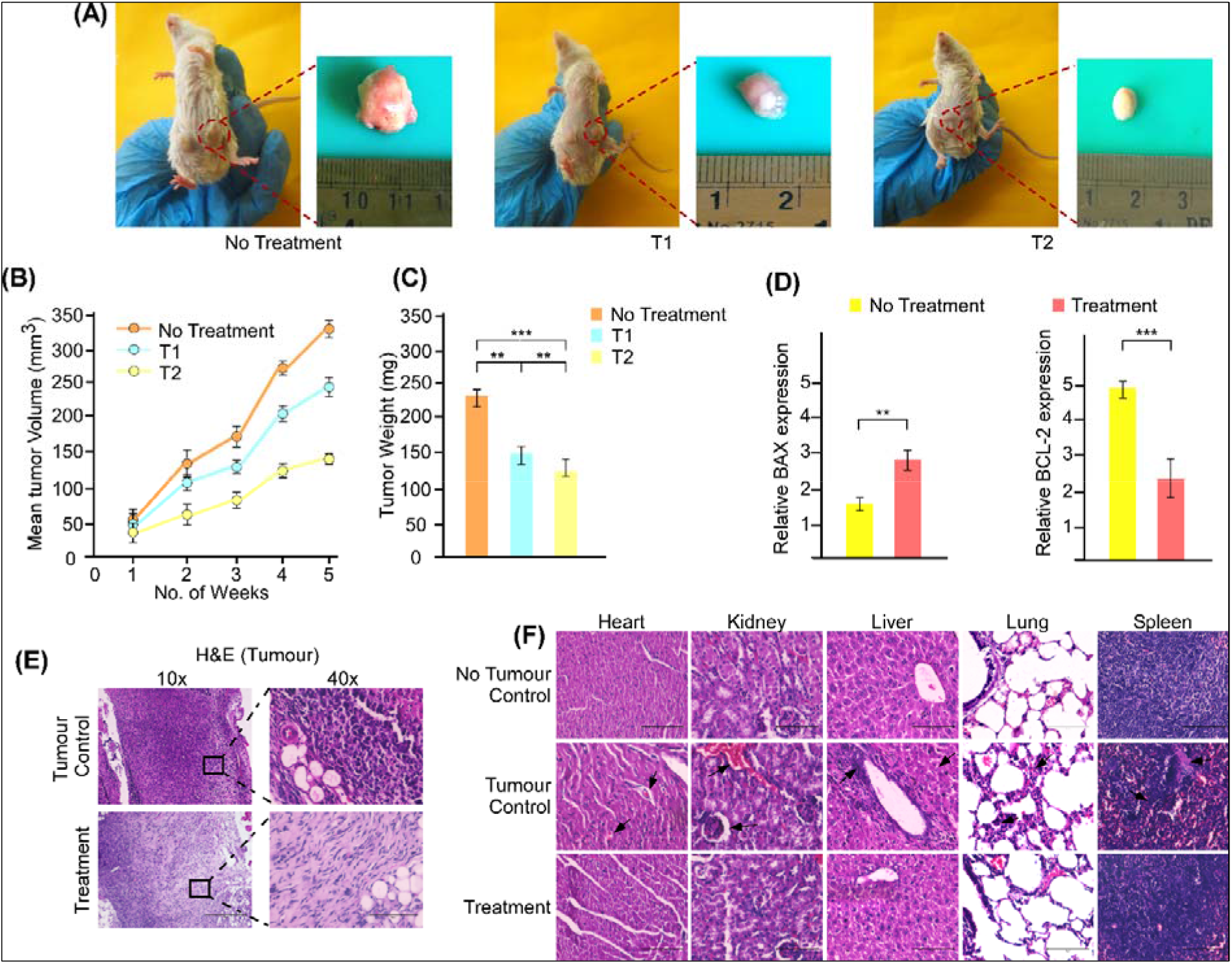
In-vivo experimentations to authenticate anti-cancerous potential of BSO: [(A) Left Panel] Images depicting the various sized tumours (formed as a result of 4T1 induction) excised from the BalB/c mice without treatment, [(A) Middle Panel] treated with T1 & [(A) Right Panel] T2 in order to highlight the efficacy of BSO against the tumor formed. T1 represents 20 mg/kg while T2 represents 30 mg/kg body weight. Graphical representation to exhibit the modifications in the (B) Mean tumor volume as well as (C) the tumor weight. The following set of experimentations were performed with the T2 dosage. (D) Graphical representation denoting the changes in the expression pattern of Bax & Bcl-2 by means of qRT-PCR. (E) Eosin haematoxylin staining was employed to histopathologically analyse the excised 4T1 tumors from both (Upper Panel) tumor control & (Lower Panel) treated sets followed by phase contrast imaging. (F) Histological sections of heart, kidney, liver, lung & spleen from these mice were stained with haematoxylin and counter-stained with eosin and their microscopic analysis was performed for histopathological examinations of toxicity in tissue. Values are mean ± SEM of three independent experiments in each case or representative of typical experiment. **P <0.01, and ***P <0.001.

### Analysis of tumor induced peripheral toxicity and the effect of BSO on the same

To assess peripheral toxicity imposed by tumor development and effect of BSO on the same, various parameters were evaluated. The activities of ALT, AST and ALP are indicative of metabolic health and any changes in the activities of these enzymes particularly point towards hepatic toxicity (Hall et al., 2012) (Hall and Cash, 2012). The data obtained showed that tumor development resulted in significantly increased activities of ALT, AST and ALP [Fig. 10A (ii), (iii) & (iv)] while treatment with BSO was able to restore near normalcy activity levels of these enzymes. Further, we also observed that tumor bearing mice had significantly elevated activity of LDH compared to Control and treatment with BSO reduced serum LDH activity thus, establishing its efficacy as an anti-tumor agent [Fig. 10A (i)].

**Figure 10.**
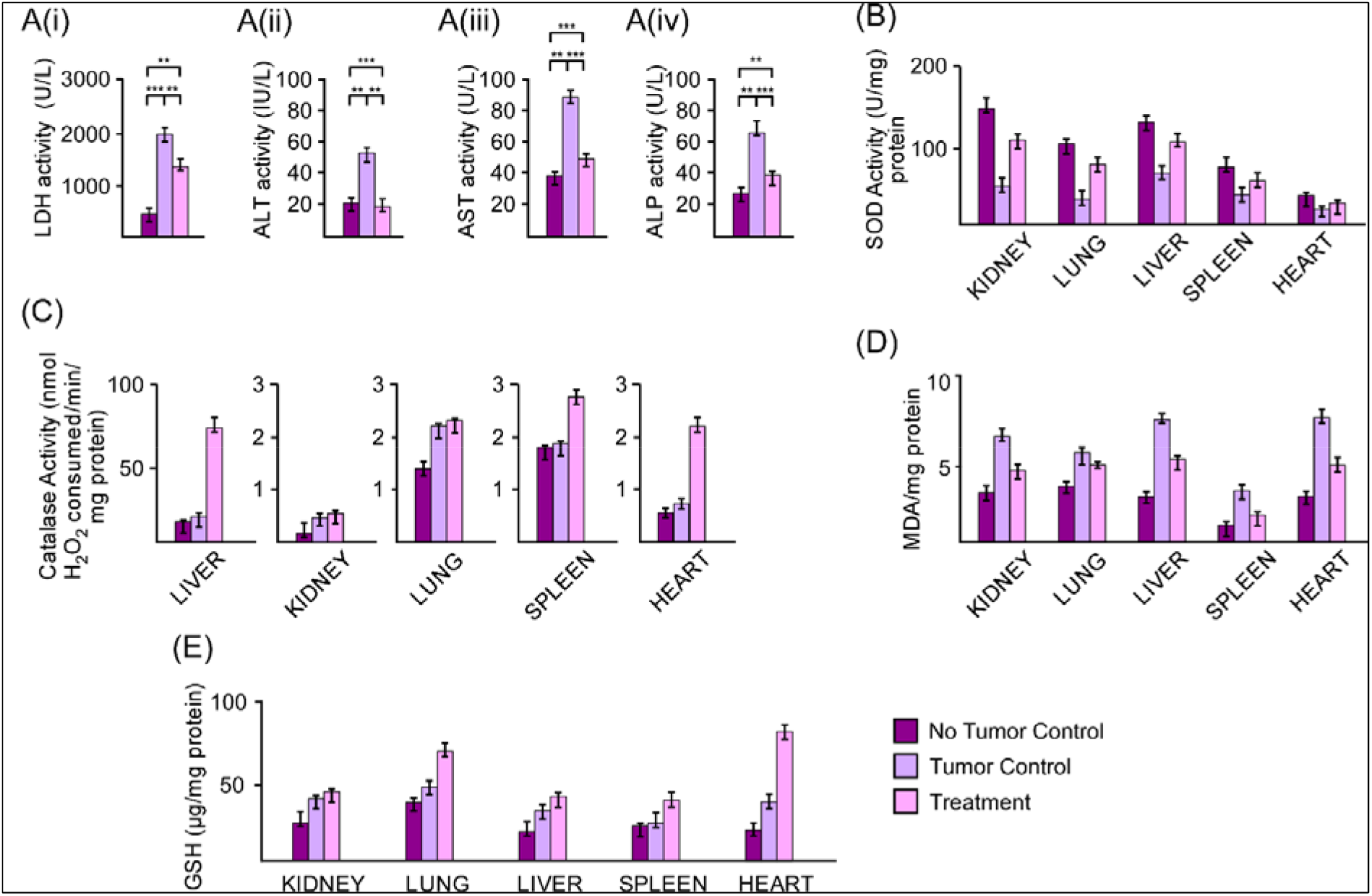
Inspection of in-vivo Serum toxicity & BSO clearance from living system: (A-E) Blood was collected from the sacrificed BalB/c mice to monitor any systemic changes resulting from oral gavaging of BSO. (A) Graphical Representation, portraying the Serum levels of (i) LDH, (ii) ALT, (iii) AST and (iv) ALP gives an idea of the Serum toxicity. The extent of (B) SOD activity, (C) Catalase activity, (D) Malonaldehyde & (E) GSH were also presented graphically for the organs collected from the sacrificed BalB/c mice of both Untreated & Treated Sets. Values are mean ± SEM of three independent experiments in each case or representative of typical experiment. **P <0.01, and ***P <0.001.

**Figure 11.**
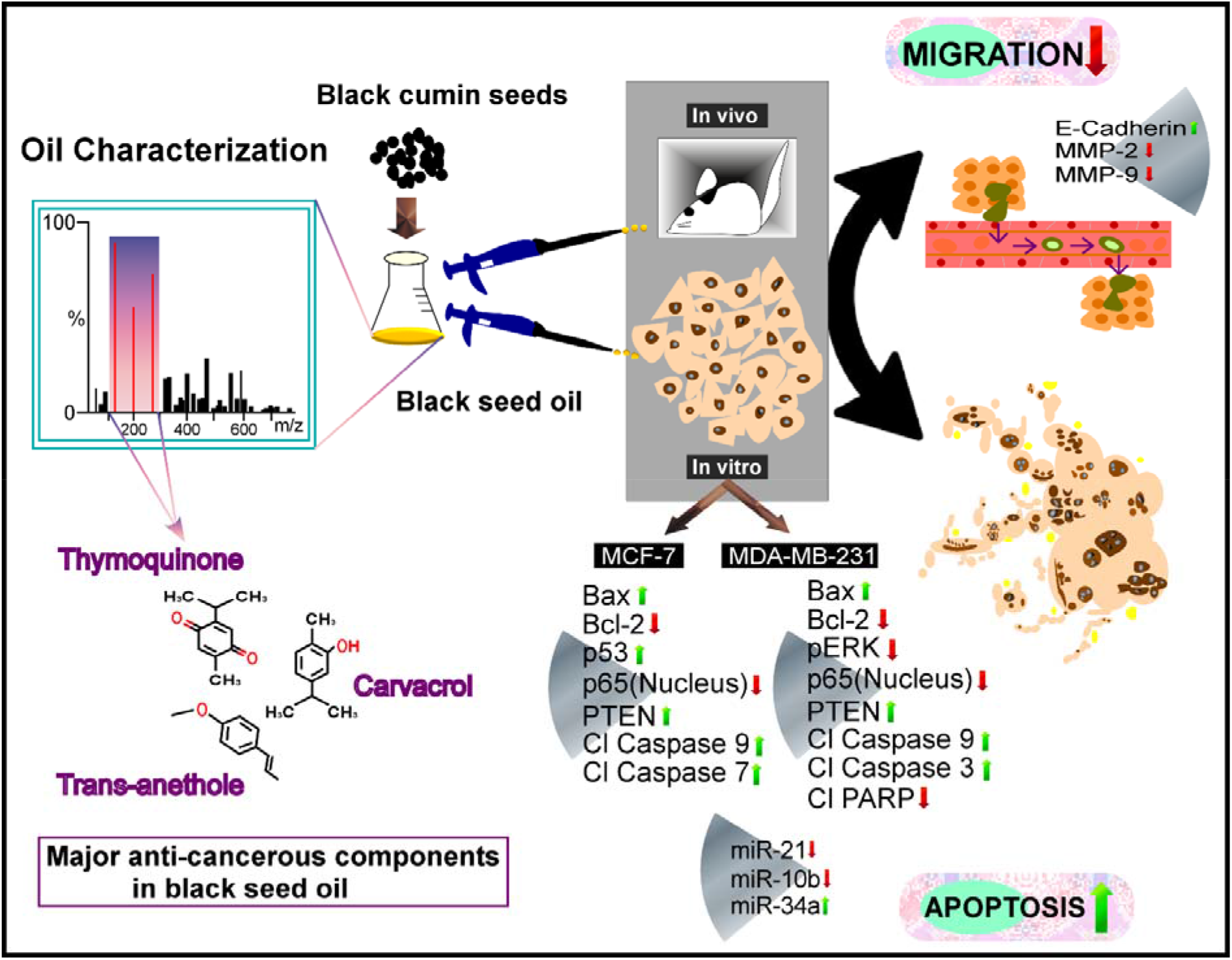
Graphical Abstract. Schematic illustration, portraying the anti-migratory & apoptotic role of BSO in breast Cancer

A physiological state oxidative stress is closely associated with tumor development (Reuter et al., 2010). In concurrence with this, estimation of lipid peroxidation levels from the 10% PMS of heart, liver, lungs, kidneys and spleen of experimental animals showed a significant increase in the lipid peroxidation in the tumor bearing mice as compared to control which was lowered upon BSO treatment[Fig.10(D)]. Also, the ratio of SOD to Catalase was much higher for tumour bearing mice. Treatment with BSO resulted in a decrease SOD to Catalase proportion thus indicating that the BSO treatment was countering the physiological oxidative stress due to breast tumour and re-establishing the anti-oxidant defences [Fig.10(B & C)]. This was further corroborated by the fact the tumour-induced reduction in GSH levels were restored upon BSO treatment [Fig. 10(E)].

## DISCUSSION

The upcoming era of new age healthcare welcomes phytochemicals & their irreplaceable contribution in the medical field owing to their availability, low toxicity & scarce side effects (Choudhari et al., 2020). Our employment of Black seeds in this study is to extensively evaluate the anti-cancerous effect of the oil it houses in totality in comparison to its commercially available, supremely processed active components like Thymoquinone, Trans-Anethole & Carvacrol. With this notion in mind, we have commenced this study with the pharmacognostical analysis of the crude black seed samples. In the process of preparing the active oil (detailed structure documented in the material & methods part), we have extracted the required ingredients of the seeds by using organic solvents n-Hexane, Di-Ethyl Ether & Dichloromethane in a multistep extraction.

The evaluation of its properties via HRMS & GC-MS (Fig. 3 & 4) validated the presence of the active components in Black seed oil. As previously reported, the different active components, TQ, TA & Carvacrol individually have anti-cancerous effect (Ahmad et al., 2019). So the current study aimed to utilise their synergistic value in curbing breast cancer growth and migration at a concentration so that it does not harm the normal cells. GC-MS analysis also justified this observation as it clearly portrayed the percentage of TQ, TA & Carvacrol present in the formulation was much lower as compared to the amount housed by the purified components which is in sync with our observation in the performed Cytotoxicity Assay, wherein the BSO is capable of causing cancer cell death likewise the individual ingredients while bringing about minimal to no toxicity to the PBMC population. Generation of ROS has been considered to be a major determinant in the activation of the mitochondrial death cascade (Bae et al., 2011). The increasing level of ROS in the treated cells is also conclusive of the enhanced oxidative stress which tackles tumorigenesis in case of both the MCF-7 & MDA-MB-231 [Fig. 6(C)]. The SEM micrographs, furthermore confirmed the ongoing apoptotic changes triggered via caspase activation in the cells under treatment exhibiting signs of nuclear fragmentation, blebbing & chromatin condensation [Fig. 6(D-F)].

Another deadliest hallmark of cancer includes its distant metastasis which makes treatment avenues more limited and difficult to handle (Seyfried and Huysentruyt, 2013). Interestingly, BSO successfully halted the migratory nature of the breast cancer cells as evident from our Wound Healing & Transwell Assay [Fig. 7 (A-D)]. As the cytoskeleton remains one of the biggest pinnacle of targets for any anti-migratory drug, we closely looked into the lamellipodia and filopodia assembly forming the pioneers of cell-cell contact (Arjonen et al., 2011; Machesky, 2008) & observed considerable reduction in them which can be correlated to the inefficient migration of the BSO treated cells [Fig. 7 (E)]. As the said assembly plays a pivotal role in invasion & metastasis, the compromise achieved by BSO indeed gives it a promising score. To further validate the anti-migratory flair of BSO, we observed the changes in expression of E-Cadherin, MMP-2 & MMP-9 at the translational level [Fig. 7 (F)]. Our observation of E-cadherin upregulation solidified the identification of BSO to limit migration in both the breast cancer cell lines. In addition, substantial downregulation of MMP-2 & MMP-9 expression also aimed at the similar conclusion.

In order to investigate deep into the molecular basis of the anti-migratory & apoptotic potential of BSO, we evaluated an array of proteins & micro RNAs (Fig. 7) which has not been unveiled so far. Here, an enhanced expression of p53 in case of treated MCF-7 cells along with an elevated Bax expression in both the cell lines is consistent with its identity as a facilitator of apoptotic cell death. As already reported, Bax works in consortium with another protein, Bcl-2 & their balance plays a very prominent role in Apoptosis. In this study as well, we found reduction in the Bcl-2 expression in both cell lines on treatment with BSO. Next, we studied another tumour suppressor, the phosphatase and tensin homolog deleted from chromosome 10 (PTEN) that has been postulated earlier to negatively regulate PI3K/AKT signaling which affects cellular integrity, thereby prompting apoptosis (Chalhoub and Baker, 2009). Our study observed an enhanced PTEN expression when treated, in both MCF-7 & MDA-MB-231. This study also showed a depletion in poly (ADP-ribose) polymerase (PARP)& decline in p65 nuclear expression in BSO treated cells, contributing the modifications to the reduced capacity of PARP-1 in revitalizing the DNA damage caused by BSO in the treated sets & inactivation of anti-apoptotic target genes such as Bcl-2 family proteins y in both the cell lines. Moreover, BSO treatment brought down the expression of pERK kinase in MDA-MB-231 cells, critical to its role in tumor invasion and metastasis. Delving more into the mechanism by which BSO brings about the apoptotic cellular demise, we explored the expression changes in the Caspases. We observed a substantial enhancement in their expression which along with the above observations confirmed the apoptotic demise of the breast cancer cells. Additionally, while we studied expression of micro RNAs (Fig. 8E), we found reduction in the expression of miR-21 that brings about sufficient upgradation in Bcl-2 expression. Similarly, another tumour inducer, playing havoc in cancer metastasis is miR-10b. Its enhanced expression is indicative of several hallmarks of cancer - increased metastasis, enhanced invasive capability both *in vitro* and *in vivo*. On the contrary, we also studied a potent tumour suppressor, miR 34a that showed a significant upregulation owing to its capacity to lower the expression of Bcl-2, an antiapoptotic protein.

Further, to confirm the effects of BSO in minimising solid tumours & restrain migration in mammary cancer cells, we opted to investigate in a detailed *in vivo* study wherein we observed a successful tumour regression in the treated sets (Fig. 9A). Besides, the near normalcy reinstated in the tumour sections on H & E staining from the ample morphological discrepancies in the Control sets is further concordant with the action of BSO against breast cancer (Fig 9 E). In addition, the detailed histopathological study of the different organ samples excised upon the sacrifice of the animal reassured the similar. Moreover, tumour induced peripheral toxicity studies were conducted which further brought into limelight the safety index of BSO usage to mitigate migration & emerge instrumental in bringing about apoptosis.

Amassing the above observations serve the perfect staging of BSO as a formidable phytochemical entity which as a whole encompasses to bring about a fair relief in the sphere of breast cancer research.

## CONCLUSION

The present study accounted for a generalised better cytotoxic credibility of the volatile oil against breast cancer compared to its major integrants-Thymoquinone, Trans-anethole & Carvacrol. Upregulation of PTEN, and Bax accompanied reduced expression of Bcl2 & nuclear p65 on treatment with oil. Additionally, the results directed a reduction in expression of oncomiRs-miR-21, miR-10 & significant upgradation of miR-34a on administering the oil as a whole. Summarising, this study for the first time highlights a cost effective method for isolation of black seed oil and evaluates the cumulative effect of the integrants working in harmony against breast cancer.

## Supporting information

Supplementary Section S1

## ABBREVIATIONS

BSO: Black Seed Oil
Cyt C: cytochrome C
DAPI: 4’,6-diamidino-2-phenylindole
FACS: fluorescence-activated cell sorting
MTT: 3-(4,5-dimethylthiazol-2-yl)2,5-diphenyltetrazolium bromide
ROS: Reactive Oxygen Species
SEM: Scanning electron microscope
TA: Trans-Anethole
TNBC: Triple negative breast cancer
TQ: Thymoquinone

## ACKNOWLEDGEMENT

We would like to sincerely thank Dr. Anirban Roy, Dr. Manisha Ahir & Dr. Saurav Bhattacharya for proof reading the article. We would like to thank Dr. Rajesh Bolleddu & Mr. Manajit Bora for helping with the Pharmacognostical data. Gratitude is due to Ms. Sneha Mitra (CU-BD-CoE) & Mr. Prothyush Sengupta for their technical support.

## Funding

This work was supported by the Ministry of AYUSH (sanction no: Z.28015/29/ 2016-HPC (EMR)-AYUSH-A) Govt. of India.

## ETHICAL STATEMENT

The mice for the in-vivo experimentations were maintained according to the guidelines of the Institutional Animal Ethical Committee (IAEC) & approved by the same IAEC-/V/P/SC-05/2019 dated 07.08.19.

## AUTHOR CONTRIBUTIONS

SD performed the experimentations, lead data acquisition with considerable support from AG and PD who helped with the active oil extraction as well. SS & PU aided in the molecular screening protocols alongside SD while PG aided in the in-vivo work load along with MB. SG collaborated in drafting the article with SD besides participating in reviewing of the manuscript. SC helped in supervising the in-vivo experimentations while AA was in charge of the conceptualization, editing & overall reviewing of the manuscript.

## CONFLICT OF INTEREST

The author(s) declare no competing interests associated with this publication and there has been no significant financial support for this work that could have influenced its outcome.

## MATERIALS & METHODS

### Other Reagents

Carvacrol, Trans-Anethole and Thymoquinone (TQ) were purchased from Sigma Chemical Company (USA).Culture media Dulbecco’s Modified Eagle’s Medium (DMEM), L-15 medium, penicillin, streptomycin, gentamycin, L-glutamine, non-essential amino acids(NEM), amphotericin B, 3-(4,5-Dimethylthiazol-2-yl)-2,5 diphenyltetrazolium bromide, a tetrazole MTT, were procured from HIMEDIA (Mumbai, India).Fetal bovine serum (FBS) was purchased from Invitrogen (Carlsbad,USA). MTT, acridine orange, was obtained from Sigma Chemical Co (USA). Flow cytometry was performed using AnnexinV-fluorescein isothiocyanate (AnnexinV-FITC)/propidium iodide (PI) kit (BD Pharmingen, San Diego, CA). Antibodies against p53, p65, pERK, PARP, PTEN, BAX, Bcl2, E-Cadherin, MMP-2 & MMP-9 were obtained from Santa Cruz Biotechnology (Santa Cruz, USA). Hoechest 33342, Phalloidin, 40, 6-diamidino-2-phenylindole (DAPI) were purchased from Invitrogen (Carlsbad, USA).

### Cell line & Cell culture

Human breast cancer cell lines MCF-7 & MDA-MB 231 were purchased from National Centre for Cell Science (Pune, India). The cells were cultured in 10% FBS, penicillin (100μg/mL), streptomycin (100μg/mL), and gentamycin (100μg/mL) containing complete DMEM in a 37°C humidified atmosphere of 5% CO_2_ incubator and MDA-MB 231 was maintained in L-15 media supplemented with aforementioned components but in absence of CO_2_. Care was taken to perform the cell culture experiments in biosafety cabinet under sterile condition. Human whole blood was collected from adult healthy volunteers with prior consent in heparinized vacutainer blood collection tubes (BD, Franklin Lakes, NJ). 100 mL Whole blood was diluted with 150 mL of RPMI-1640. It was then layered in centrifuge tubes onto 120 mL of Histopaque-1077 gradient and centrifuged the peripheral blood mononuclear cells (PBMC) layer. Then PBMC layer was washed twice with PBS and cells were re-suspended followed by culture in RPMI-1640 with 10% FBS.

### Cytotoxicity assay

Determination of the toxic potential of Black Seed Oil (BSO) was done with the aid of MTT (Sigma) to find out the percentage cell survival on treatment with BSO. The MCF-7 & MDA-MB 231 cells were harvested, counted and transferred to 96-well plates and incubated for 24hrs prior to the addition of BSO. The oil was administered at increasing concentrations from 5-50 μg/mL and the treated cells were incubated for another 24hrs. MTT (3-[4, 5-dimethylthiazol-2-yl] - 2, 5- diphenyltetrazolium bromide) (5mg) was dissolved in 1 mL of phosphate-buffered saline (PBS), and 25 μL of the MTT solution was added to each of the 96 wells. The plates were wrapped in aluminium foil and incubated at 37°C for 4hrs. The solution in each well, containing media, unbound MTT and dead cells, was removed by suction, and 200 mL of DMSO was added to each well. The plates were then shaken, and the optical density was measured using a microplate reader; Multiskan™ GO Microplate Spectrophotometer at 575 nm. The Control set was considered to have a cell viability of 100%.

### Deciphering Cellular Apoptosis by Flow Cytometry

In order to determine the apoptotic dose of BSO on MCF-7 &MDA-MB 231 cells, apoptosis assay was performed. Cells positive for apoptosis and necrosis was measured by the Annexin V/propidium iodide assay. The externalization of phosphatidyl serine as a marker of earlystage apoptosis was detected by its ligand Annexin-V protein conjugated to FITC, whereas membrane damage due to late-stage apoptosis/necrosis was detected by the binding of PI to nuclear DNA. The continuously cultured breast cancer cells were harvested in 6-well plates and incubated for 24hrs. Then various doses of BSO were applied on them and the treated cells were incubated for another 24hrs. There were control cells which remained untreated. Next day, cells were washed in binding buffer (10mM HEPES, pH 7.4; 140mM NaCl; 2.5mM CaCl2) and incubated in the dark for 10 min at room temperature in 100 ul binding buffer containing Annexin V-FITC (40 ul/ml) and PI (1ug/ml). After incubation, 400 ul binding buffer was added to each sample and cells were kept on ice. The 488 nm laser was used for excitation and FITC was detected in FL-1 by a 525/30 BP filter while PI was detected in FL-2 by a 575/30 BP filter. Standard compensation was done in the Quanta SC MPL Analysis software (Beckman Coulter) using single-stained and unstained cells. For each sample, 20 000 cellswere analyzed and apoptotic (Annexin V+, PI-), necrotic (Annexin V+, PI+) and live (Annexin V-, PI-) cells were expressed as percentages of the 20 000 cells.

### Measurement of intracellular ROS

Sets of 2 * 10^6^ cells (approximately) were treated by BSO at a specific dosage for 24 h. At the end of incubation, the cells were scrapped and pellet collected by centrifugation at 300 g for 5 min at Room Temperature. The pellets were collected and suspended in 1 mL of PBS, pre-warmed to 37°C. 2 ml of H_2_DCFDA working solution was then added from stock to make the final concentration of 2 mM. The cell suspensions were incubated for 20 min at 37 C and was protected from light. After that, FACS analyses of the samples were carried out with FACSVerse at an excitation at 488 nm and emission at 520 nm utilising the FACSuite software.

### Cellular morphology examination by Scanning Electron Microscopy (SEM)

The morphology of breast carcinoma cells under different conditions were observed under Scanning Electron Microscope (SEM). By following the protocol of Bhattacharya *et al*., 2015 the cells were grown on poly-_L_-Lysine (1:10) coated glass slide (1 cm^2^) overnight at 37 C in 5% CO_2_. Next day, cells were treated with specific dose of BSO and kept in the CO_2_ incubator for overnight. Then, the cells were washed by sodium cacodylate buffer, and fixed with 2.5% glutaraldehyde buffered in sodium cacodylate for 1 h. The samples were washed with sodium cacodylate buffer thrice, dehydrated through a series of alcohol concentrations (10%, 30%, 50%, 70%, 90% and 100%), and subjected to air drying. The samples were then visualized by EVO-18 special edition SEM (ZEISS, Germany).

### Wound healing assay

Cancer cell migration under different conditions was examined by bidirectional wound healing assay. Both the MCF-7 & MDA-MB-231 cells were grown in 12 well plates & at a confluency of around 80 to 90%, a linear scratch created in the cell monolayer formed a bidirectional wound. Migration was quantitated by a semi-automated, computer-assisted process with aid of a person blinded with respect to the experimental treatment. The data from triplicate wells were calculated as the means ± S.E.M. The rate of migration in case of control cells was taken as 100% and healing rate of other plates were compared with respect to control cells.

### Transwell migration assay

Transwell migration assay was performed using cell culture inserts (BD Biosciences, Sparks, USA) with pores (8 mm). Placing the inserts in the 12-well sterile cell culture plates, 2*10^5^ cells were seeded in them. DMEM (10% FBS) was then added to the lower chamber of the 12-well Plate & left overnight. Dose addition was done the following day & after another 24hrs, the non-migrated cells in the upper compartment was carefully washed away. The cells in the insert were then washed, fixed with 3.7% formaldehyde and then permeabilized with 100% methanol and at last, stained with Giemsa for 30min. Cells present in the lower part were counted and the number of migratory cells was expressed as an average number of cells per microscopic field.

### Western blot analysis

MCF-7 & MDA-MB-231 cells treated with BSO were harvested and lysed using RIPA buffer (Sigma, USA) supplemented with protease and phosphatase inhibitor cocktail (Cell Signaling Technologies, Beverly, MA). Protein concentrations were measured using Bicinchoninic acid (BCA) assay Kit (Merck, Germany). An equal amount of protein was loaded and separated by SDS-PAGE, which was then transferred to polyvinylidene fluoride membranes (Millipore, Germany) and incubated with primary antibodies - anti-β actin, p53, p65, PARP, pTEN, pERK, Bcl-2, Bax, E-Cadherin, MMP-2 & MMP-9 (purchased from Santa Cruz Biotechnologies, USA). HRP conjugated secondary antibodies (Sigma, USA) was then added followed by developing using chemiluminescence. The ratio of the intensity of the target protein to that of the β-actin loading control was calculated to represent the expression level of the protein under investigation.

### RNA isolation and quantitative real-time PCR

Total RNA isolation was carried out by Phenol Chloroform method using TRIZOL Reagent after verifying the strength and purity of total RNAs with the help of ND1000 spectrophotometer (Thermo) followed by gel-ectrophoresis on urea denaturation gel. 1 ug of total RNAs was reverse transcribed using Transcriptor First Strand cDNA Synthesis Kit and oligo (dT)18 primer (Roche, Switzerland & Biobharti, India). In case of miRNA detection, the miR-specific stem-loop primers were used along with same cDNA synthesis kit. qRT-PCR was performed with 50 mg sample using Light cycler 96 (Roche) and SYBR green ready mix (Roche) and the mentioned primers (Supplementary Table 1). U6 and 18s were used as internal controls for miRNA and mRNA quantification, respectively. Gene and miR expression was defined from the threshold cycle (C_q_), and relative expression levels were calculated by using the 2^DCq^ method after normalization with reference gene.

The primer sequences used for the following:

**Supplementary Table 1:**
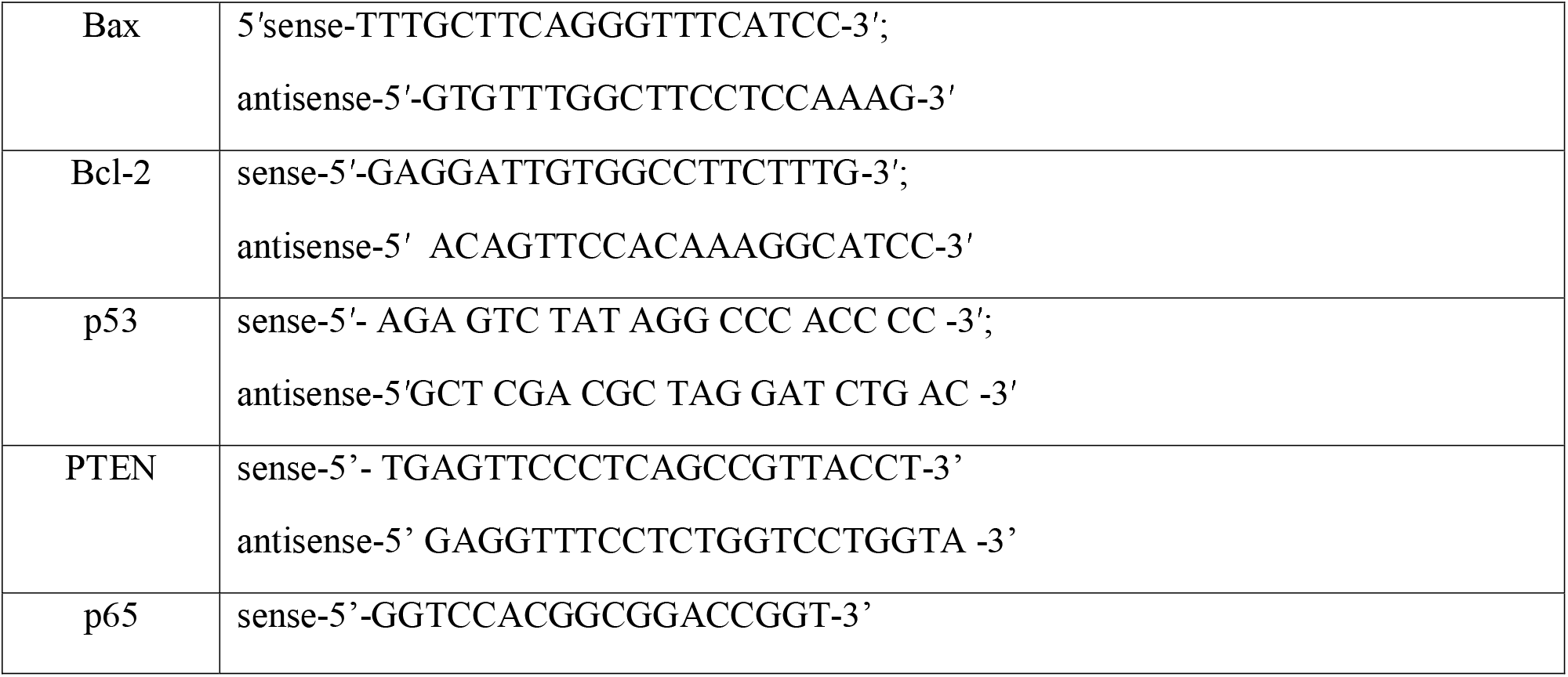
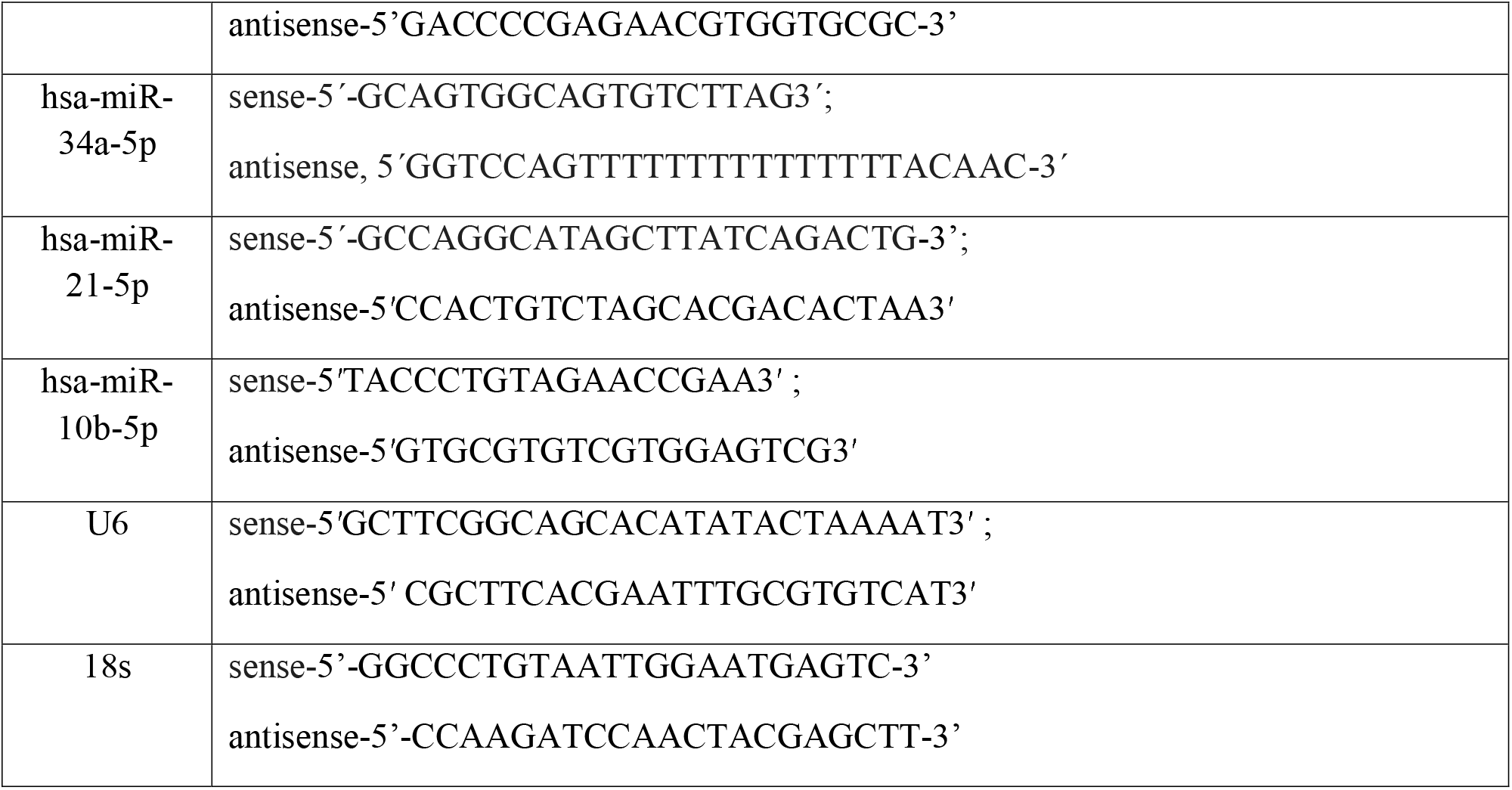
PRIMER SEQUENCES USED IN THE STUDY

### Serum toxicity assay

Blood was collected from the retro-orbital plexus of the treated mice post-anaesthetization. The isolated blood was incubated at 4°C in a slant position for 1 hour. Thereafter, clot was removed by centrifugation at 2000 g for 15 minutes and serum was collected. The activities of Aspartate transaminase (AST; EC 2.6.1.1), Alanine transaminase (ALT; EC 2.6.1.2), Lactate dehydrogenase (LDH; EC 1.1.1.27) and Alkaline phosphatase (ALP; EC 3.1.3.1) were estimated using standard kits as per manufacturer protocol. Assay kits for LDH and ALP were procured from Reckon Diagnostics P. LTD. (India), and assay kits for AST and ALT were purchased from ROBONIK®(India).

### Preparation of Post-Mitochondrial Supernatant (PMS)

A 10% w/v homogenate of the harvested organs was made using chilled phosphate buffer (0.1 M, pH ~ 7.4) containing KCl (1.17% w/v). The nuclear debris was separated by centrifuging the homogenate at 800 g for 5 minutes at 4°C. The supernatant was again centrifuged at 10, 800 g for 20 minutes at 4°C and the pellet was discarded. The supernatant thus obtained was the 10% PMS which was used to assess oxidative stress parameters (Kaur et al., 2006). Estimation of biochemical parameters associated with oxidative stress: Catalase (CAT; EC 1.11.1.6) activity in the 10% PMS from different tissues was estimated by measuring Catalase mediated decomposition kinetics of H_2_O_2_ into H_2_O and O_2_ using a spectrophotometer (λ = 240 nm) (Claiborne 1985).Superoxide dismutase (SOD; EC 1.15.1.1) activity was estimated based on the principle of inhibition of pyragallol auto-oxidation by SOD enzyme in sample according to the protocol described by Marklund, 1985 (Marklund 1985). Lipid peroxidation in the tissue 10% PMS was measured using thiobarbituric acid which forms a chromogenic adduct with malonaldehyde, the final product of lipid peroxidation (Wright et al., 1981). Reduced glutathione (GSH) levels in the tissue 10% PMS was estimated using the DTNB (5,5’-dithiobis-(2-nitrobenzoic acid) reagent as per previously described protocol (Jollow et al., 1975).

### Histopathological studies

Mice of the treated groups were sacrificed and the heart, liver, kidney, spleen, lungs and tumour tissues were resected. The tissues were fixed in 10% formalin, were subjected to graded dehydration and paraffin embedding as per standard protocols. The paraffin embedded tissues were cut into 5-7 μm slices and stained with hematoxylin and eosin or used for IHC. Microscopic examination was done at 10X (Scale Bar: 450 μm) or 40X (Scale Bar 100 μm) magnifications using an EVOS XL Cell Imaging System, Thermo Fisher Scientific (Upadhyay et al., 2019).

## Notes

### Competing Interest Statement

The authors have declared no competing interest.

